# *Hypanus brevis*: a newly resurrected Eastern South Pacific stingray lineage revealed by integrative taxonomy

**DOI:** 10.64898/2026.02.25.708098

**Authors:** Alan Marín, Fabiola Zavalaga, Renato Gozzer-Wuest, Luis E. Santos-Rojas, Lorenzo E. Reyes-Flores, Ruben Alfaro, Philippe Béarez, Eliana Zelada-Mázmela

**Affiliations:** Laboratorio de Genética, Fisiología y Reproducción, Facultad de Ciencias, Universidad Nacional del Santa, Chimbote, Perú; Área Funcional de Investigaciones en Biodiversidad, Instituto del Mar del Perú - IMARPE, Callao, Perú; Innovations for Ocean Action Foundation (I4OA), Lima, Peru; Ecobiotech Lab S.A.C. Trujillo, Perú; UMR 7209 BioArch, CNRS-MNHN, Muséum national d’Histoire naturelle, CP 56, 43 rue Buffon, 75005 Paris, France

**Keywords:** Cryptic species, marine conservation, molecular phylogeny, Myliobatiformes, taxonomic resurrection

## Abstract

*Hypanus brevis* (Garman, 1880) and *Hypanus dipterurus* (Jordan & Gilbert, 1880) are currently considered as a conspecific lineage of the Eastern Pacific “diamond stingray”. This group has been the subject of nomenclatural disputes for about 145 years, with *H. brevis* accepted as the junior synonym. To clarify the historical confusion surrounding this lineage, we employed an integrative taxonomic approach using specimens from the Eastern North and Eastern South Pacific (ENP and ESP). Both single (COI) and multilocus (COI - 16S rRNA) genetic analyses revealed a distinct evolutionary unit in the ESP. While morphological analyses detected subtle differences between ENP and ESP specimens (e.g., disc length and internostril width), most characters exhibited significant overlap, suggesting low evolutionary divergence. A Bayesian molecular clock analysis estimated this divergence at approximately 5.6 Ma. In accordance with Garman’s (1880) original description based on specimens from Paita (northern Peru), we formally resurrect *H. brevis* from synonymy with *H. dipterurus*. Our findings suggest an anti-tropical speciation pathway, with core populations of *H. brevis* and *H. dipterurus* restricted to the temperate waters of the ESP and ENP, respectively. Notably, a single, fixed COI haplotype was detected in all *H. brevis* specimens from the north-central Peruvian coast. This result may indicate a severe bottleneck event, raising concerns about the genetic health and long-term viability of this vulnerable species. Finally, we analyzed historical fishery data of *H. brevis* to infer its current population status, suggesting targeted conservation measures and precautionary management to prevent further loss of genetic diversity.

## 1. INTRODUCTION

Species of the genus *Hypanus* Rafinesque, 1818 (Myliobatiformes: Dasyatidae) are characterized by their wide pectoral fins that are fused to the head, a prominent row of thorns on nape accompanied by one or more thorns on shoulder in adults, and a distinct but low ventral skin fold on tail (Last et al. 2016a). Most *Hypanus* species hold significant commercial value throughout their distribution range in the Atlantic and Pacific Oceans. They are either targeted by artisanal fisheries or frequently retained as valuable bycatch (Feitosa et al. 2021; Campos-León et al. 2025; Mejía-Falla et al. 2025). Due to their high market demand, unmanaged fishing practices, and their k-selected life history traits (e.g., slow growth and low fecundity) (Yamaguchi et al. 2021), half of the species in this genus are currently listed as threatened on The IUCN Red List of Threatened Species (IUCN 2025).

The genus *Hypanus* was considered a junior synonym of *Dasyatis*, until it was resurrected by Last et al. (2016b). It is now composed of eleven species (Fricke et al. 2025), including seven species from the Western Atlantic: *Hypanus americanus* (Hildebrand & Schroeder, 1928), *H. berthalutzae* Petean, Naylor & Lima, 2020, *H. geijskesi* (Boeseman, 1948), *H. guttatus* (Bloch & Schneider, 1801), *H. marianae* (Gomes, Rosa & Gadig, 2000), *H. sabinus* (Lesueur, 1824), and *H. say* (Lesueur, 1817); one species from the Eastern Atlantic: *Hypanus rudis* (Günther, 1870); and three species from the Eastern Pacific: *Hypanus dipterurus* (Jordan & Gilbert, 1880), *H. longus* (Garman, 1880), and *H. rubioi* Mejía-Falla, Navia, Cardeñosa & Tavera, 2025. However, a recent phylomitogenomic analysis suggested the existence of three additional evolutionary units within this genus (*H.* aff*. americanus*, *H.* aff*. guttatus*, and *H.* aff. *say*), warranting the need for a thorough taxonomic revision (Petean et al. 2024).

In the Eastern Pacific, the diamond stingray, *H. dipterurus*, is found from southern California (USA) to northern Chile (Lamilla et al. 1995; Almendras et al. 2025). This species is currently listed as Vulnerable on The IUCN Red List of Threatened Species, primarily due to intense and largely unmanaged fisheries across most of its distribution range (Pollom et al. 2020). The diamond stingray is one of the most captured species by Mexican artisanal fisheries (Simental-Anguiano et al. 2022). In Colombia and Ecuador, its capture is limited to bycatch (Calle-Morán and Béarez 2020; Puentes et al. 2022), while in Peru, artisanal fisheries target this species (González-Pestana et al. 2022; Campos-León et al. 2025). Despite its intense exploitation, high demand, and signals of population declines (Pollom et al. 2020), no formal management plans have been established in Peruvian stingray fisheries yet (González-Pestana et al. 2022). Furthermore, a long-standing taxonomic confusion with a junior synonym, *Dasyatis brevis* (Garman, 1880), may have skewed historical catch estimates.

The diamond stingray has faced a long period of confused nomenclature history, due to two independent and nearly simultaneous descriptions published in 1880, that led to a confusing synonymy (Miller et al. 2014). The first description was published in May 1880 by Jordan and Gilbert, who named it *Dasybatis dipterurus* based on specimens collected from San Diego Bay in California (Jordan and Gilbert 1880). Then, in October of the same year, Garman described *Trygon brevis*, based on two specimens from Paita, Peru (Garman 1880). Garman (1913: 396) synonymized the two species (*D. dipterurus* and *T. brevis*) under the name *Dasybatus brevis*, considering wrongly that the date of publication of Jordan and Gilbert’s work was 1881, thus giving priority to his own description from 1880. This inappropriate synonymization has led to long-standing ongoing confusion, dividing the opinion of modern researchers (Nishida and Nakaya 1990; Nelson et al. 2004; Miller et al. 2014) until Eschmeyer (1998) recognized the nomenclatural priority of *D. dipterurus* Jordan & Gilbert, 1880 over *T. brevis* Garman, 1880. Ebert (2003), followed by Love et al. (2005) and Mundy (2005) then reclassified the latter as a junior synonym of *Dasyatis dipterura*, and finally Last et al. (2016b) reassigned the species to the genus *Hypanus* (currently *Hypanus dipterurus*). Despite these taxonomic clarification efforts, the status of *H. dipterurus* and *Dasyatis brevis* remains controversial, even 145 years after their initial descriptions.

Inaccurate taxonomic revisions and hidden diversity can lead to a cascade of errors resulting in dire conservation consequences. This is particularly true for commercially exploited batoids, due to their high vulnerability to overfishing (White and Last 2012; Hosegood et al. 2020). Treating multiple distinct species as a single group, or inversely, a single species managed as different units, can mask localized population declines and thus threaten the sustainability of these batoids (Siskey et al. 2019; Chatzispyrou and Koutsikopoulos 2023). Furthermore, assessments of conservation status rely on the analysis of fisheries statistics from landing reports and geographical distributions based on taxonomic revisions (IUCN 2024). The use of flawed catch statistics and imprecise distribution ranges resulting from taxonomic uncertainties can lead to inaccurate conservation status assessments (Marshall et al. 2009).

DNA sequence analysis represents a powerful tool for achieving taxonomic clarity, particularly in groups with complex morphological identification, such as batoids, where overlapping traits challenge the use of traditional methods (Sales et al. 2019; White et al. 2022; Marín 2025; Cunha et al. 2026). Genetic studies have played a crucial role in uncovering cryptic diversity within morphologically conservative ray species, leading to significant formal taxonomic revisions (Last et al. 2016b; White et al. 2022). For example, a phylogenetic analysis of the southern stingray, *Hypanus americanus*, revealed that it was paraphyletic (Petean et al. 2020). This finding led to the discovery of a new species, the Lutz’s stingray, *H. berthalutzae* by Petean et al. (2020). Additionally, genetic studies have led to the reinstatement of names that were previously treated as junior synonyms. These include the Indo-Pacific spotted eagle ray, *Aetobatus ocellatus* (Kuhl, 1823), which was previously considered a junior synonym of *A. narinari* (Euphrasen, 1790) (White et al. 2010); *Neotrygon trigonoides* (Castelnau, 1873), previously synonymized with *N. kuhlii* (Müller & Henle, 1841) (Borsa et al. 2013); and *Zearaja brevicaudata* (Marini, 1933) (currently *Dipterurus brevicaudatus*), which was confused with *Zearaja chilensis* (Guichenot, 1848) (currently *D. chilensis*) (Gabbanelli et al. 2018), among others.

Most taxonomic and phylogenetic revisions including *H. dipterurus* samples have primarily focused on populations from the Eastern North Pacific (hereafter referred to as ENP) (e.g., Naylor et al. 2012; Last et al. 2016b; Petean et al. 2020). On the other hand, the populations of *H. dipterurus* from the Eastern South Pacific (hereafter referred to as ESP) have not yet been assessed. Consequently, there is a significant knowledge gap regarding the connectivity and gene flow among the populations from the northern and southern hemispheres. Furthermore, a recent study suggested that *H. dipterurus* from the ESP may represent a distinct evolutionary lineage (Petean et al. 2024). This study aims to evaluate the evolutionary history and species boundaries of the Pacific diamond stingray, *H. dipterurus*. First, we generated DNA sequences from *Hypanus* specimens collected in the ESP and compared them with available conspecific sequences from the ENP populations. Then, we provided a redescription of *H. dipterurus* from the ESP populations based on an integrative taxonomic approach, which led to the formal resurrection of *H. brevis* (Garman, 1880). Lastly, we evaluated historical and contemporary landing trends and discussed conservation priorities and management strategies for sustainable *Hypanus* fisheries in the ESP.

## 2. MATERIALS AND METHODS

### Ethical statement

All stingray samples used in this study were obtained from Peruvian artisanal fisheries, landings, wholesale markets, local retail markets, and from activities conducted aboard research vessels, from January 2017 to January 2026. Consequently, no animals were sacrificed specifically for the purposes of this study, and no special permissions were required as the target species are not under protection.

### Collection of voucher specimens

Four whole specimens of ESP *H. dipterurus* were collected and deposited in the Scientific Marine Collection of the Peruvian Marine Research Institute (IMARPE) under the catalog numbers IMARPE-020476 to IMARPE-020479. These include a juvenile caught (IMARPE-020476) as bycatch off Redondo Beach in Miraflores district, collected at the landing site “*Muelle de Pescadores*” in Chorrillos district (Peru). Three additional commercial specimens (IMARPE-020477 to IMARPE-020479), caught at San José Beach in Lambayeque (Peru), were bought in a wholesale fish market located in Chiclayo city (Lambayeque, Peru) (see Table 1 and Fig. 1). A small piece of tissue was excised from the oral epithelium of the buccal cavity of each sampled specimen using a sterile scalpel and preserved in ethanol 96 % for further DNA analysis.

**Fig. 1.**
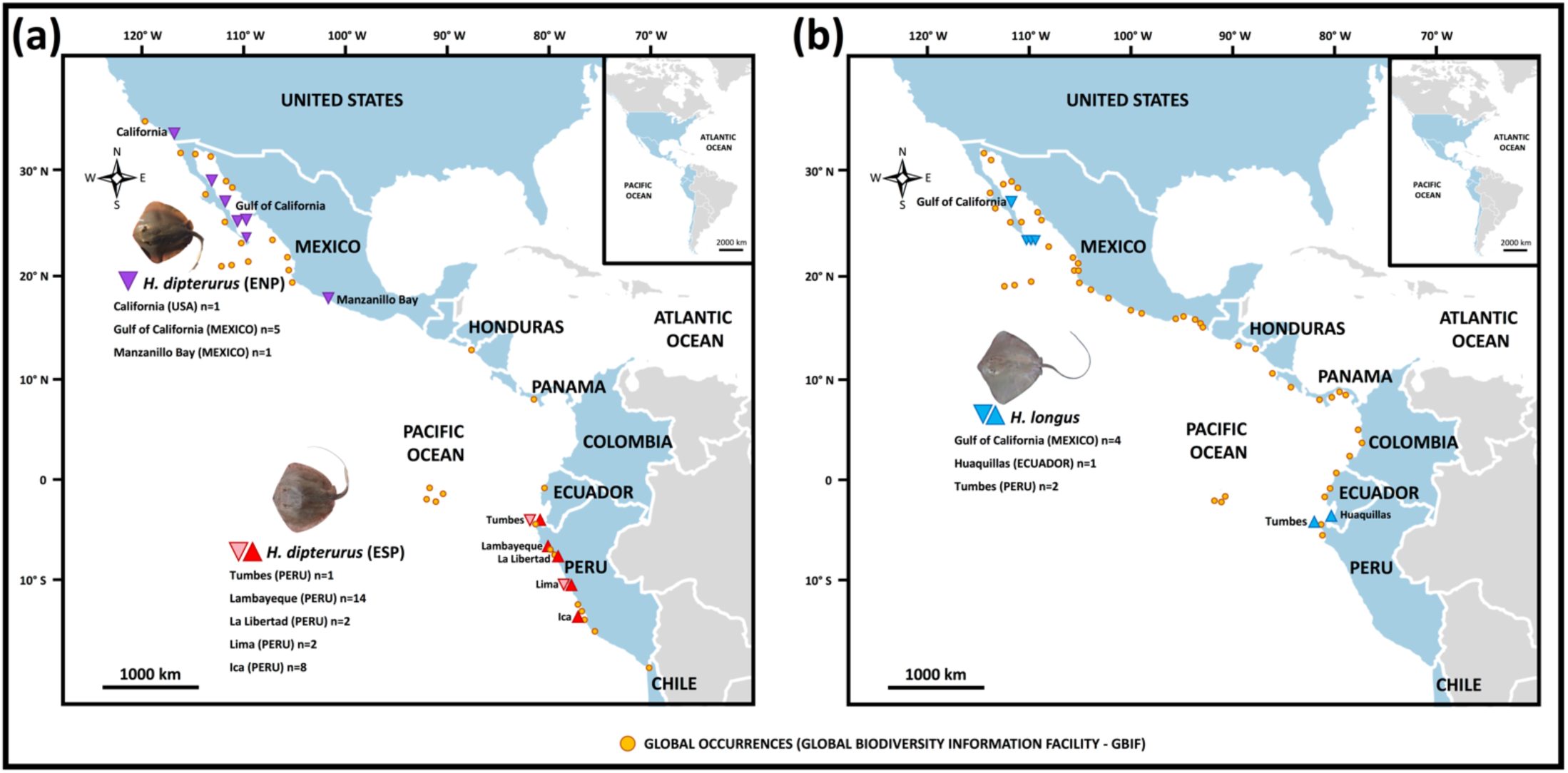
Sampling sites of *Hypanus* stingrays from the Eastern Pacific coast. Panel (a): Inverted purple triangles pinpoint sampling locations of *H. dipterurus* from the Eastern North Pacific (ENP) population with DNA sequences available in GenBank. Red triangles pinpoint sampling locations of *H. dipterurus* from the Eastern South Pacific (ESP) population collected in this study. Inverted pink triangles denote sampling sites of *H. dipterurus* samples whose DNA sequences were retrieved from the GenBank and BOLD databases. Panel (b): Inverted blue triangles indicate sampling locations of *H. longus* from the ENP population whose DNA sequences were retrieved from GenBank. Upright blue triangles indicate the sampling locations of *H. longus* from the ESP population collected in this study. Yellow circles represent global occurrences for *H. dipterurus* (a) and *H. longus* (b) as reported in the Global Biodiversity Information Facility (GBIF) database.

**Table 1.**
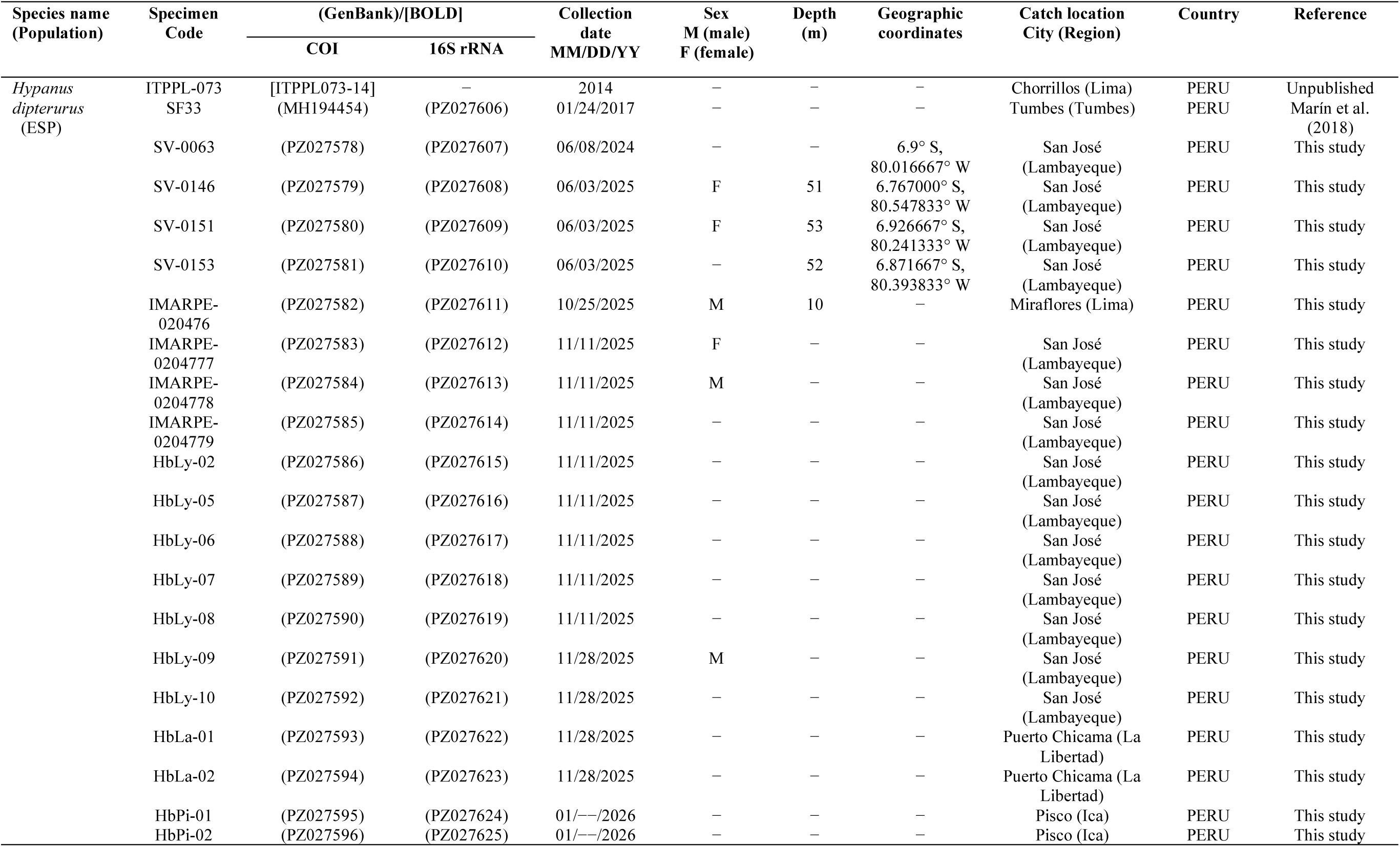
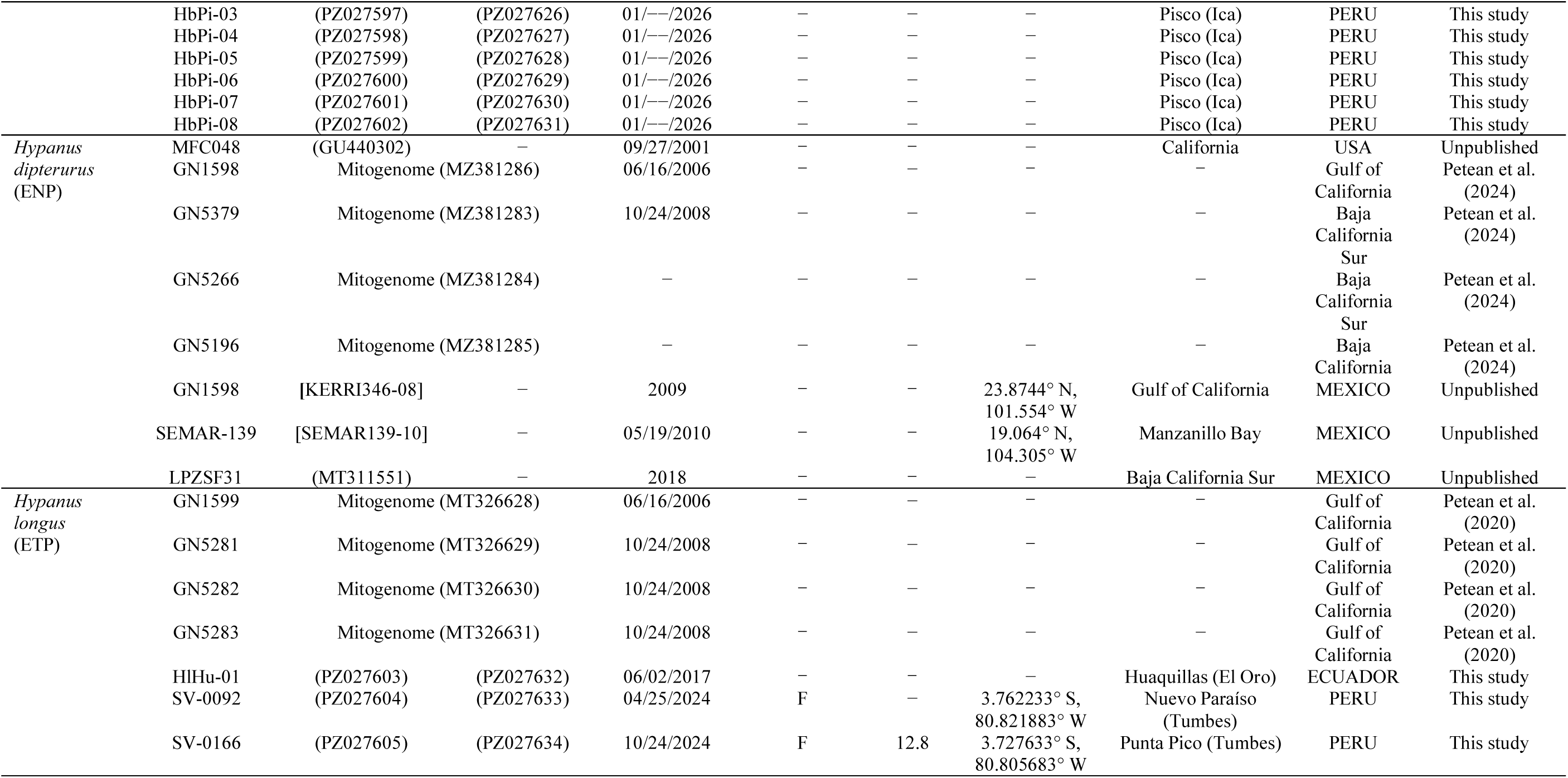
List of samples analyzed in this study. GenBank and BOLD accessions are written between parentheses and brackets, resp Eastern North Pacific; ETP: Eastern Tropical Pacific.

### Tissue sample collection

We also collected additional tissue samples from whole and incomplete specimens of ESP *H. dipterurus* (n = 22) and *H. longus* (n = 3) sourced from commercial fisheries, landings, and a wholesale fish market in the Lambayeque, Ica, and Tumbes regions (Peru; sample code prefixes HbLy, HbPi, and SF in Table 1). Tissue sample from one *H. longus* individual were obtained from a local market in Huaquillas city (Ecuador, sample code prefix HlHu in Table 1). Muscle samples (ESP *H. dipterurus* and *H. longus*) were also obtained during IMARPE research activities aboard research vessels and labeled with the prefix SV (without voucher specimens) (see Table 1 and Fig. 6a-b). Small pieces of muscle from fillets, fins, and torsos were collected and preserved in commercial ethanol 96 % in 1.5 mL Eppendorf microtubes and stored at-15 °C until DNA extraction.

### Molecular identification and data analyses DNA isolation

Total genomic DNA was isolated using the cetyltrimethylammonium bromide (CTAB) protocol described by Zuccarello and Lokhorst (2005) with two modifications. In the organic extraction phase, the original chloroform:isoamyl alcohol (24:1) mixture was replaced with a phenol:chloroform:isoamyl alcohol in a final ratio of 25:24:1. During the subsequent DNA precipitation step, an additional 1/10 volume of sodium acetate (3 M, pH 5.2) was added to enhance precipitation efficiency. DNA quantification was estimated in an Epoch spectrophotometer (BioTek Instruments, Winooski, VT, USA). The genomic DNA was diluted to a final concentration of 50 ng/µL and used in PCR amplifications.

### PCR amplification

A partial fragment (707 bp) of the mitochondrial COI gene was obtained using the universal fish primer set FishF1 (5’-TCAACCAACCACAAAGACATTGGCAC-3’) and FishR1 (5’-TAGACTTCTGGGTGGCCAAAGAATCA-3’) (Ward et al. 2005). For the amplification of the mitochondrial 16S rRNA gene fragment (632 bp), we used the universal primers 16Sar-L (5’-CGCCTGTTTATCAAAAACAT-3’) and 16Sbr-H (5’-CCGGTCTGAACTCAGATCACGT-3’) (Palumbi et al. 1991). All PCR products were amplified in a total reaction volume of 10 µL with the following master mix composition: 0.1 µL of *Taq* polymerase 5 U/µL (HiMedia, MBT060, Maharashtra, India), 0.9 µL of 10x PCR Buffer, 0.76 µL of MgCl_2_ (25 mM), 0.5 µL of dNTPs (2.5 mM), 0.1 µL each primer (50 µM), 1 µL of genomic DNA (50 ng/µL), and 6.54 µL of ultrapure H_2_O. Two different PCR amplification conditions were used, depending on the DNA marker. For the COI marker, it consisted of an initial denaturation for 5 min at 95 °C, followed for 35 cycles of denaturation step 30 s at 94 °C, annealing for 40 s optimized from a range of 54-60 °C, and elongation for 60 s at 72 °C. A final extension step was performed for 10 min at 72 °C. For the 16S rRNA, the initial denaturation was 5 min at 95 °C, followed for 35 cycles of denaturation for 45 s at 94 °C, annealing for 45 s at 58 °C, and elongation for 90 s at 72 °C.

The final extension lasted 5 min at 72 °C. Successful PCR amplifications were verified in a 1.5 % agarose gel electrophoresis.

### DNA sequencing

Prior to DNA sequencing, PCR products were purified using FastAP Thermosensitive Alkaline Phosphatase and Exonuclease I (Thermo Fisher Scientific, USA). Purified amplicons were sequenced bi-directionally using the same primer sets and the BigDye Terminator v3.1 Cycle Sequencing Kit (Applied Biosystems, Waltham, Massachusetts, USA) on a ABI PRISM 3500 Genetic Analyzer (Applied Biosystems, Hitachi, Foster City, CA, USA). All sequencing reactions were performed at the Laboratory of Genetics, Physiology, and Reproduction of the National University of Santa. Electropherograms were visually revised using MEGA 7 (Kumar et al. 2016), consensus sequences were obtained by aligning forward and reverse complemented sequences, and primer positions were removed.

### Estimation of genetic diversity and population differentiation

We used Arlequin v3.5 (Excoffier and Lischer 2010) to quantify genetic variation within each analyzed *H. dipterurus* population (ENP and ESP) and to estimate diversity metrics. These include haplotype number, haplotype diversity “*Hd*”, nucleotide diversity “*π*”. The relationships among haplotypes resulting from the COI and COI + 16S rRNA datasets were assessed using Median-joining networks in HaplowebMaker tool (Spöri and Flot 2020). To assess deviations from selective neutrality and infer historical demographic events, we calculated Fu’s *F_S_* and Tajima’s *D* in both datasets (COI and COI + 16S rRNA) using Arlequin. Significant negative values of these indices are indicative of recent population expansion, whereas positive values suggest population bottlenecks or balancing selection. Statistical significance was evaluated with 10,000 coalescent simulations, with threshold significance levels set at P < 0.02 (*F_S_*) and P < 0.05 (*D*).

To quantify the degree of genetic divergence between the *H. dipterurus* samples collected from ENP and ESP populations, we used the Analysis of Molecular Variance (AMOVA) implemented in Arlequin. Pairwise fixation indices (Φ*_CT_*, Φ*_SC_*, and Φ*_ST_*) were estimated using the Kimura 2-parameter model (K2P) (Kimura 1980) for the COI and concatenated COI + 16S rRNA datasets. Accordingly, total variance was grouped into the following components: Φ*_CT_*: variation among groups, Φ*_SC_*: variation among populations within groups, and Φ*_ST_*: variation among all populations. Two populations were defined: 1) the ENP population, comprising 8 DNA sequence samples sourced from GenBank, whose geographic origin indicated California in the USA, and Gulf of California and Manzanillo Bay in Mexico; 2) the ESP population, represented by 27 DNA sequences of which 25 sequences were determined in this study (Ica region n = 8, Lambayeque region n = 14, La Libertad region n = 2, and Lima region n = 1, see Table 1). Two additional Peruvian sequences were sourced from GenBank (Tumbes region n = 1) and BOLD (Lima region n = 1) databases.

### *_D_*Bayesian phylogenetic analysis

The phylogenetic relationships among the DNA sequences of *Hypanus* species were inferred using two independent Bayesian analyses based on single COI locus dataset and a concatenated COI + 16S rRNA multilocus dataset. The partitioned Bayesian approach on the multilocus dataset was conducted to enhance the phylogenetic signal of the 16S rRNA gene and improve the reliability of species delimitation (Nylander et al. 2004). Although a preliminary single-locus analysis of the 16S rRNA gene recovered ESP *H. dipterurus* and ENP *H. dipterurus* as separate clades, the relatively low genetic distance (K2P: 1.7 %) indicates a recent divergence between these two clades. Such shallow genetic divergences are often associated with limited phylogenetic signal (Philippe et al. 2011). This may result in weakly supported topology or overly confident statistical nodal support (Chen et al. 2017).

The single (COI) and multilocus (COI + 16S rRNA) datasets consisted of 55 and 50 sequences, respectively. These included 25 sequences for ESP *H. dipterurus* (Peru) and three sequences for *H. longus* (Ecuador and Peru) determined in this work. They were analyzed along with all available conspecific sequences from Mexico and the USA that were retrieved from BOLD and GenBank databases. Additionally, four sequences of *H. say* (sister taxon of *H. dipterurus*) and *H. americanus* (sister taxon of *H. longus*), and two sequences of *H. sabinus* (sister taxon to all *Hypanus* species) (Rosenberger 2001; Petean et al. 2024; Mejía-Falla et al. 2025), were retrieved from the databases and included in the analyses. Two bluespotted stingrays (*Neotrygon indica* Pavan-Kumar, Kumar, Pitale, Shen & Borsa, 2018 and *N. kuhlii*, subfamily Neotrygoninae) and the Daisy stingray (*Fontitrygon margarita* (Günther, 1870), subfamily Urogymninae) were used as outgroups. DNA sequences from the COI and 16S rRNA genes were aligned independently and trimmed using MEGA 7. Pairwise genetic distances were calculated using the K2P model (Kimura 1980), as implemented in MEGA 7. We assessed signals of nucleotide saturation in the protein-coding COI dataset using DAMBE 7 (Xia 2018). Next, the best-fit model of evolution for each aligned matrix were assessed with jModelTest2 (Darriba et al. 2012) under the corrected Bayesian Information Criterion (BICc). The best evolutionary model for the COI and 16S rRNA gene datasets were HKY + G and HKY + I, respectively.

The Bayesian phylogenetic analyses were conducted in MrBayes 3.2.7 (Ronquist et al. 2012), as implemented on the CIPRES Science Gateway 3.3 server (Miller et al. 2010). The analyses were conducted with the data partitioned by gene and codon position. In both datasets, the COI and the concatenated COI + 16S rRNA, the setup parameters consisted of two independent runs of four Markov chains each, conducted for 10,000,000 generations. Trees were sampled every 1,000 generations and the first 25 % of the sampled trees were discarded as burn-in. Quality check of output files consisted on an Average Standard Deviation of Split Frequencies (ASDSF) below 0.01 and Effective Sample Sizes (ESS) for all parameters exceeding 200. The ESS were verified in Tracer v1.7.2 (Rambaut et al. 2018). The consensus trees were visualized with FigTree version 1.4.4. (http://tree.bio.ed.ac.uk/software/figtree/).

### Calibrated molecular dating analysis

To estimate the divergence ages among target *Hypanus* species, we conducted a relaxed molecular dating analysis using the concatenated multilocus dataset (COI + 16S rRNA: 1,116 bp). The dataset was partitioned into three subsets: COI first and second codon positions, COI third codon position, and the 16S rRNA gene. Since the single and multilocus phylogenies consistently recovered the ENP and ESP *H. dipterurus* populations as distinct monophyletic clades, we applied the more general substitution model GTR + I + G to all partitions.

To test each divergence node and identify where constraints were necessary to maintain stability, we performed initial exploratory sensitivity analyses without topological constraints to independently. This allowed us to implement two independent minimalist constraint models to avoid over-parameterization while enabling independent topology inference: 1) a single-anchored model and 2) a double-anchored model.

1) The single-anchored model: this model used the base of the Dasyatidae clade as a deep calibration point, based on the divergence between the lineages of *Dasyatis* and *Urogymnus* estimated by Aschliman et al. (2012) using fossil and molecular data. The node age was calibrated using an “offsetlognormal” distribution, which effectively incorporates uncertainty by providing a hard minimum age while allowing for a probabilistic tail toward older dates (Ho and Phillips 2009). Specifically, the node was assigned a lower bound of 57.4 million years ago (Ma) via offsetlognormal(57.4, 64.9, 0.5). Additionally, a structural constraint was enforced *a priori* to maintain the sister relationship between Neotrygoninae and Dasyatinae and to prevent chronological distortions. This constraint was necessary because, while standard Bayesian inference (non-clock) recovered the expected topology (see “Bayesian phylogenetic analysis” in Results section), preliminary molecular clock tests recovered Neotrygoninae as sister to Urogymninae. This discrepancy was probably caused by the additional parameters and branch-length constraints imposed by the molecular clock model, which failed to resolve deep basal relationships
2) The double-anchored model: this alternative approach employed the previously mentioned prior parameters of the single-anchored model, with the addition of an internal calibration point based on the final closure of the Isthmus of Panama. For this purpose, the node representing the divergence between the Atlantic *H. say* and ENP *H. dipterurus* was assigned an offsetlognormal distribution(3.1, 3.5, 0.4), based on the final closure of the Isthmus of Panama approximately at 3.1-3.5 Ma (O’dea et al. 2016). The results of this double-anchored sensitivity analysis were used to assess the robustness of our single-anchored model.

Finally, we applied a lognormal(-5.2, 1.0) prior to the mean clock rate to model the typical slow mitochondrial mutation rate (0.2-0.3 % per million years) documented for chondrichthyan taxa (Martin et al. 1992). The molecular clock analysis was conducted using the same MCMC computational framework and quality check of output files as the phylogenetic inference described in the previous subsection. However, to accommodate the increased complexity of the relaxed clock model (IGR), the MCMC run was increased to 50,000,000 generations to reach stationary.

### Morphometric and meristic analysis

Morphometric and meristic analyses were conducted on four complete, fresh specimens of *Hypanus* (Table 1), following primarily Petean et al. (2020), with additional measurements taken from Jordan and Gilbert (1880). A total of 53 morphometric measurements and three meristic counts were recorded for each specimen. The complete list of measurements and counts, including abbreviations and definitions, is provided in Table 5.

All body measurements were taken in a straight line using a caliper with a precision of 0.1 mm; measurements exceeding 170 mm were taken using dividers and read with a steel ruler with a precision of 1 mm. After measurements, specimens were fixed in 10 % formalin, rinsed, preserved in 70 % ethanol, and deposited in the IMARPE Scientific Collection under the catalog numbers IMARPE-020476, IMARPE-020477, IMARPE-020478, and IMARPE-020479. Comparative morphometric data were obtained from the original descriptions of *Trygon brevis* by Garman (1880), considering the two syntypes under the collection number MCZ Ichthyology S-422, one female and one male from Paita, Peru; and from Lamilla et al. (1995). We compared with morphometric data of two syntypes of *Dasybatis dipterurus* from Jordan and Gilbert (1880) description; and with those given by Beebe and Tee-Van (1941) and Nishida and Nakaya (1990). Measurements are expressed in millimeters (mm) and in percentages (%) of the disc width (% DW). In some cases, measurements originally expressed in inches were converted to millimeters. When necessary, measurements were also standardized from proportions of total length (TL) or disc length (DL) to ensure comparability. Taxonomic validity and nomenclature follow Fricke et al. (2025), while collection codes are based on Sabaj (2020). All specimens from the IMARPE fish collection were also used for species identification through genetic analyses.

### Fishery data analysis

A database of chondrichthyan landings from 1996 to 2024, geolocated by fishing zones, was provided by the IMARPE, in response to a request for access to public information under Peruvian transparency provisions. In total, the database comprised 182,282 landing records, classified by geolocation (latitude and longitude), fishing gear, common and scientific names, year, and landed volume. Records of batoid species, particularly those classified as *Dasyatis brevis* and the generic scientific designation *Hypanus* sp., were filtered to describe small-scale fishing activity targeting species under analysis. Consequently, both labels were extracted for descriptive purposes. Additionally, fishing areas of the small-scale fisheries were classified into four zones by latitude: Tropical North, North, Center, and South, based on IMARPE’s classification available in the Peruvian Atlas of Small-Scale Fisheries (Marín-Soto et al. 2017). Additionally, the distance to the coast of the reported catches was estimated using the Fenix R Studio package (Marín-Abanto 2018). The data were used to estimate trends in landings and to show the marine areas within the Peruvian Sea with the highest occurrence of this species’ fishing activity. Furthermore, to discover its economic value, annual average off-vessel prices and total monetary landing values were estimated using open-access data collected and stored by IMARPE (2025). Nearly 2,108 records of nominal off-vessel prices for the research target species from 2009 to 2024 were obtained. The nominal off-vessel prices were converted from Peruvian soles (PEN) to US dollars (USD) using monthly PEN/USD exchange rates published by the Peruvian Central Bank (BCRP 2025a). Additionally, nominal prices were converted to real prices by deflating them using the consumer price index (CPI), published monthly by the BCRP from January 1991 onwards, with December 2021 as the baseline (i.e., IPC = 100) (BCRP 2025b).

## 3. RESULTS

### DNA sequencing

We successfully obtained genomic DNA, PCR products, and DNA sequences for the COI (*H. dipterurus* n = 25, *H. longus* n= 3) and 16S rRNA (*H. dipterurus* n = 26, *H. longus* n= 3) gene markers. Note that for one *H. dipterurus* individual (voucher SF33, see Table 1), only the 16S rRNA was sequenced in this study, as its COI sequence had been previously published (Marín et al. 2018) and deposited in the GenBank database under accession MH194454. After the removal of the primer sequences, the DNA sequences for the COI and 16S rRNA markers were 655 bp and 590 bp in length, respectively. All DNA sequences generated in this research were submitted to the GenBank database. COI sequences: *H. dipterurus* from PZ027578 to PZ027602; *H. longus* from PZ027603 to PZ027605. 16S rRNA sequences: *H. dipterurus* from PZ027606 to PZ027631; *H. longus* from PZ027632 to PZ027634.

### Genetic diversity indexes and neutrality tests

Genetic diversity indices for the ENP population were relatively high (*Hd* = 0.8333 - 0.8571) whereas the ESP population exhibited extremely low diversity (*Hd* = 0 - 0.1477), characterized by the fixation of a single COI haplotype across 27 specimens (Table 2). *π* followed a similar pattern, being higher in the ENP (COI: 0.003461, COI + 16S rRNA: 0.000906), and notably lower in the ESP population (COI: 0.0, COI + 16S rRNA: 0.000134) (see Table 2). Neutrality test for the ENP yielded negative values for both the Fu’s *F_S_* and Tajima’s *D*. Although these values were not significant in all cases, their negative trend deviates from expectations of a neutral population model and may reflect a recent population expansion that may be underpowered due to the limited sample size (n = 8). Neutrality test from the ESP could not be calculated for the COI gene due to the lack of polymorphism. However, the combined COI + 16S rRNA dataset, which contained two haplotypes, resulted in a non-significant Fu’s *F_S_* (-0.3167, P = 0.1648) and a Tajima’s *D* of 0.000 (P = 1.000). While these values suggest no significant departures from mutation-drift equilibrium, this is likely an artifact of the extremely low genetic diversity (e.g., one COI haplotype for 27 specimens), which limits the statistical power to detect demographic shifts. This lack of variation is consistent with a founder effect associated with a recent speciation event or a severe historical bottleneck.

**Table 2.** Summary results of genetic diversity indices and Fu’s *F s*and Tajima’s *D* neutrality tests for single locus dataset (COI) and multilocus dataset (COI + 16S rRNA) of *H. dipterurus*. ENP: Eastern North Pacific population, ESP: Eastern South Pacific population. Genetic diversity includes: number of haplotypes (h), haplotype diversity (*Hd*), and nucleotide diversity (*π*). Sample size: n. N.A.: Not Applicable.

| Population | Marker | n | h | $Hd \pm S.E.$ | $\pi$ | Fu's $F_S^*$ | Tajima's $D^{**}$ |
| --- | --- | --- | --- | --- | --- | --- | --- |
| ENP<br>(Mexico-USA) | COI | 8 | 5 | $0.8571 \pm 0.1083$ | $0.003461 \pm 0.002537$ | $-1.40322$ ($P = 0.08330$) | $-1.06428$ ($P = 0.16190$) |
| | COI +<br>16S<br>rRNA | 4 | 3 | $0.8333 \pm 0.2224$ | $0.000906 \pm 0.000897$ | $-0.88730$ ($P = 0.08860$) | $-0.70990$ ($P = 0.28510$) |
| ESP (Peru) | COI | 27 | 1 | $0.0000 \pm 0.0000$ | $0.0000 \pm 0.0000$ | $0.0000$ ($P = N.A.$) | $0.0000$ ($P = 1.0000$) |
| | COI +<br>16S<br>rRNA | 26 | 2 | $0.1477 \pm 0.0888$ | $0.000134 \pm 0.000219$ | $-0.31678$ ($P = 0.16480$) | $0.0000$ ($P = 1.0000$) |

\* $P < 0.02$ significant.
\*\* $P < 0.05$ significant

### Haplotype network analysis

The median-joining haplotype network analyses for *H. dipterurus* samples from the ENP and ESP populations are presented in Fig. 2. Our haplotype network findings from the single locus and concatenated multilocus datasets revealed a clear genetic divergence between both the ENP and ESP populations. In the COI haplotype network (top panel in Fig. 2a), five haplotypes were observed in samples from the ENP population (Mexico n = 7; USA n = 1), while a single private haplotype represented the ESP population (Peru, n = 27). The branch connecting the unique haplotype of the ESP haplogroup (H1b) to the closest haplotype from the ENP (H1d, Gulf of California) displayed a genetic distance of 18 stepwise mutations. The largest genetic distance (23 nucleotides) between both haplogroups was detected between haplotypes H1b (Peru) and H5d (Gulf of California). The haplotype network obtained using the concatenated COI + 16S rRNA dataset (bottom panel in Fig. 2a) resulted in a dominant haplotype H1b for the ESP population (Peru, n = 24), while a single satellite haplotype H2b was found in only two individuals. These two ESP haplotypes were separated by a single mutational step, represented by a deletion that was observed in the 16S rRNA gene from two specimens collected in Pisco (Ica region, GenBank accessions PZ027624 and PZ027626). The ESP haplogroup was distanced by 30 mutational steps to closest haplotype H1d in the ENP haplogroup, which comprised a total of three haplotypes from four specimens collected in Mexico. This substantial genetic distance and lack of shared haplotypes between the two haplogroups (ESP and ENP) strongly suggest that they represent isolated evolutionary units. On the other hand, the haplotype networks of both datasets (COI and COI + 16S rRNA) from *H. longus* suggest the existence of a single lineage across the Eastern Pacific (top and bottom panels in Fig. 2b, respectively). A unique dominant haplotype was detected in each population (ENP: Mexico, ESP: Ecuador and Peru), with three and five stepwise mutations between each haplotype in the single locus and multilocus datasets, respectively.

**Fig. 2.**
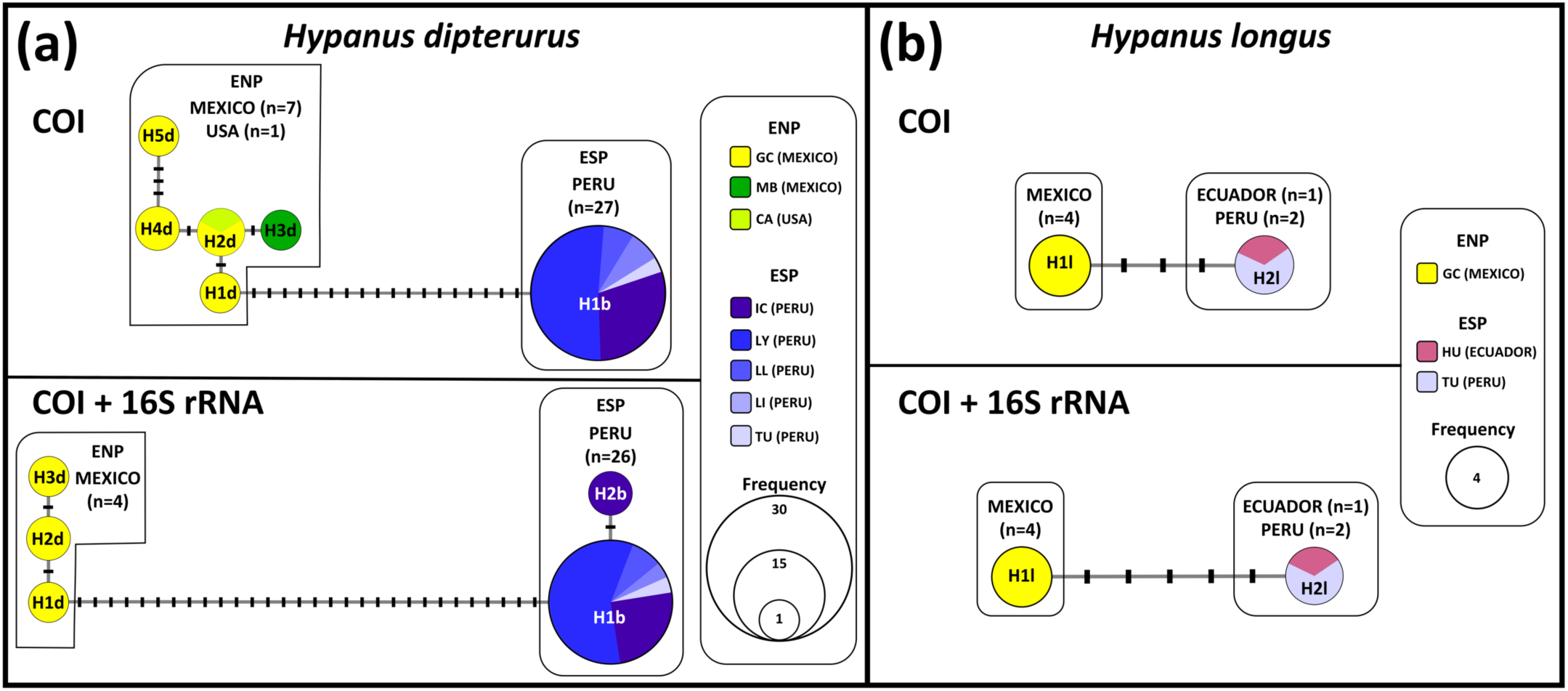
Median joining network analysis illustrating the genetic relationships and geographical distribution of two *Hypanus* lineages using DNA sequences from samples collected in the Eastern North Pacific (ENP) and Eastern South Pacific (ESP) populations. Panel (a): Haplotype network of *H. dipterurus* based on COI sequences (above) and concatenated sequences from COI + 16S rRNA (below). Panel (b): Haplotype network of *H. longus* based on COI sequences (above) and concatenated sequences from COI + 16S rRNA (below). Circles (nodes) represent unique haplotypes, where its size is proportional to the frequency of that haplotype in the sampled population. Perpendicular bars across the lines connecting haplotypes represent single mutational steps. Colors denote the geographic origin of the samples. In the ENP population: yellow color for the Gulf of California (GC) and green for Manzanillo Bay (MB), both in Mexico. Electric lime color for California (CA) in the USA. In the ESP population: raspberry pink for Huaquillas (HU) in Ecuador. Purple color for the Ica region (IC), electric blue color for the Lambayeque region (LY), neon blue for the La Libertad region (LL), pale blue for the Lima region (LI), lavender blue for the Tumbes region (TU), all located in Peru.

### Analysis of Molecular Variance (AMOVA)

The AMOVA results (Table 3) revealed high genetic differentiation between the two *H. dipterurus* populations. For the single locus (COI) dataset, the vast majority of the total variation (97.54 %) occurred between both *H. dipterurus* populations (ENP vs. ESP), while only a minor fraction (2.49 %) was observed within populations. Similarly, in the combined multilocus dataset (COI + 16S rRNA), an even higher level of variation (99.25 %) was attributed to differences between populations. Fixation indices further supported this marked divergence between ENP and ESP populations: the Φ*_CT_* values, which measure differentiation among populations, were notably high for both COI (Φ*_C T_* = 0.97, P = 0.02) and the multilocus data (Φ*_C T_* = 0.99, P = 0.16). Additionally, the consistently high and significant Φ*_ST_* values (0.98 and 0.99, P = 0.00) confirm a state of complete genetic isolation and lack of contemporary gene flow between the ENP and ESP populations. It is important to note that the increase in the P value for Φ*_C T_* observed in the multilocus dataset does not indicate a lack of differentiation, as evidenced by the near-maximum index value (0.99). Instead, this is likely an artifact of the conservative nature of the 16S rRNA gene and the limited statistical power at the among-groups level.

**Table 3.** Results of the Analysis of Molecular Variance (AMOVA) and fixation indices (Φ*_CT_*, Φ*_SC_*, and Φ*_ST_*) based on single locus (COI) and multilocus (COI + 16S rRNA) datasets from DNA sequences of *Hypanus dipterurus*. Two groups were compared: ENP (Eastern North Pacific: Mexico and USA) population group and ESP (Eastern South Pacific: Peru) population group.

| Dataset | Source of Variation | d.f. | Variance Components | % of Variation | Fixation Indices | P-value |
| --- | --- | --- | --- | --- | --- | --- |
| COI (Single) | Among groups | 1 | 9.46466 Va | 97.54 | $\Phi_{CT} = 0.97536$ | 0.02053 |
| | Among populations within groups | 6 | -0.00257 Vb | -0.03 | $\Phi_{SC} = -0.01077$ | 0.65396 |
| | Within populations | 20 | 0.24167 Vc | 2.49 | $\Phi_{ST} = 0.97510$ | 0.00000 |
| COI + 16S rRNA (Combined) | Among groups | 1 | 15.22730 Va | 99.25 | $\Phi_{CT} = 0.99249$ | 0.16618 |
| | Among populations within groups | 4 | -0.00976 Vb | -0.06 | $\Phi_{SC} = -0.08466$ | 0.36364 |
| | Within populations | 24 | 0.12500 Vc | 0.81 | $\Phi_{ST} = 0.99185$ | 0.00000 |

Fixation indices significant at $P < 0.05$

### Bayesian phylogenetic analysis

The Bayesian phylogenetic analyses of both, the single (Fig. 3a) and multilocus (Fig. 3b) datasets, further supports the conclusions drawn from the haplotype network and AMOVA findings. In both phylogenetic trees, DNA sequences from ENP and ESP *H. dipterurus* populations were recovered in two discrete monophyletic clades at the top of the trees with strong nodal support, as indicated by the Bayesian Posterior Probabilities (BPP) of 0.85 (COI) and 0.89 (COI + 16S rRNA). The sister relationship between both *H. dipterurus* populations exhibited a COI genetic distance ranging from 3.6 % (± 0.008) to 4.6 % (± 0.010), with an average value of 4 % (± 0.009) (Table S1). This distance range is substantially higher than the lowest average interspecific threshold of 0.59 % between *H. berthalutzae* and *H. rudis* (Petean et al. 2024). Our phylogenetic findings, combined with the haplotype network and AMOVA results, provide robust evidence that *H. dipterurus* populations from ENP and ESP constitute distinct evolutionary lineages.

**Fig. 3.**
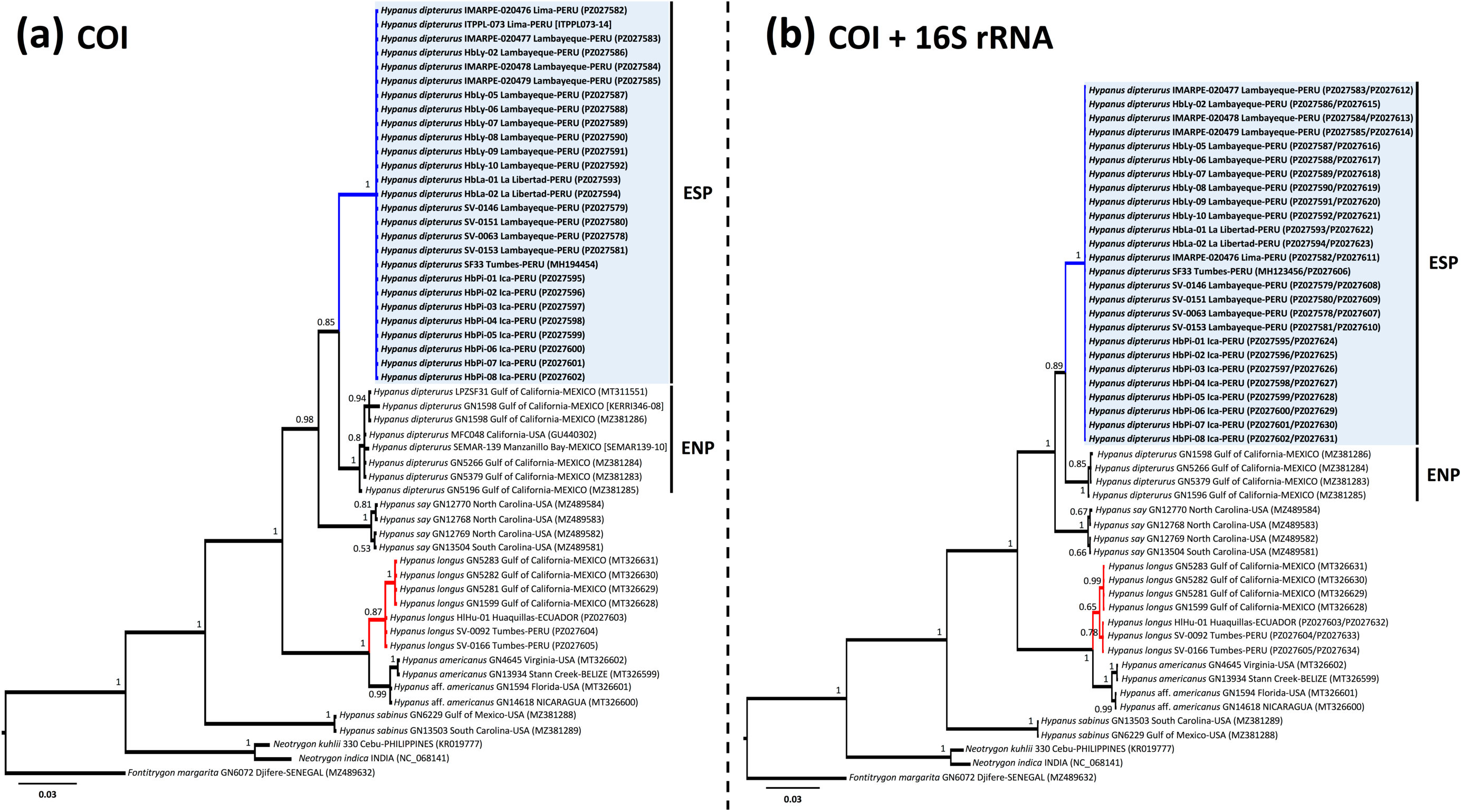
Bayesian phylogenetic trees showing the evolutionary relationships among stingrays in the genus *Hypanus*. Panel (a) show ingle locus COI marker (substitution model: HKY + G) and the tree from Panel (b) utilized the concatenated information from models: HKY + G and HKY + I). Each label shows the species scientific name, voucher name, geographical origin (when availab BOLD (between brackets) accessions numbers. The numbers next to each node indicate the Bayesian Posterior Probability. To North Pacific (ENP) and Eastern South Pacific (ESP) populations, the clade grouping all samples from the ESP population is high olor. The *H. longus* clade is highlighted in red color. *Hypanus sabinus*, the sister taxa to all *Hypanus* group, was used as an in *Neotrygon kuhlii* and *N. indica*) and the Daisy stingray (*Fontitrygon margarita*) were used as external outgroups.

Conversely, the results of the Bayesian phylogenetic analyses support the existence of a single Pacific lineage for *H. longus*. In both Bayesian phylogenetic trees, corresponding to the COI and concatenated COI + 16S rRNA dataset, all *H. longus* samples were recovered within a single monophyletic group with high statistical support (BPP = 0.87 and 0.65, respectively) (Fig. 3). They displayed low COI intraspecific genetic distances, ranging from 0 to 0.6 % (± 0.003). This maximum intraspecific value observed in *H. longus* is comparable to the minimum average pairwise interspecific COI barcoding gap (0.59 %) reported for the congeneric sister species *H. berthalutzae* and *H. rudis* (Petean et al. 2024). These findings suggest that intraspecific *H. longus* genetic variation across the Eastern Pacific does not exceed the recognized thresholds for species-level differentiation in this genus.

### Calibrated molecular clock analysis

The results of the single-and double-anchored Bayesian molecular clock analyses are presented in Fig. 4 and Fig. S1, respectively. In the single-anchored model, the most recent common ancestor (MRCA) of the Dasyatidae lineage (node a: including subfamilies Dasyatinae, Neotrygoninae, and Urogymninae) was anchored by two calibration points and placed at approximately 81.07 Ma (Upper Cretaceous; 95% HPD: 57.4 – 115.04 Ma). In contrast, the Neotrygoninae + Dasyatinae split (node b) was recovered as a topological constraint; this event occurred at approximately 70.06 Ma (95% HPD: 45.59 – 105.57 Ma), predating the K-Pg boundary. The broad 95% HPD intervals observed in these deeper nodes are consistent with previous molecular dating studies of rays (e.g. Puckridge et al. 2013; Poortvliet et al. 2015), likely related to mutational saturation of mitochondrial markers.

**Fig. 4.**
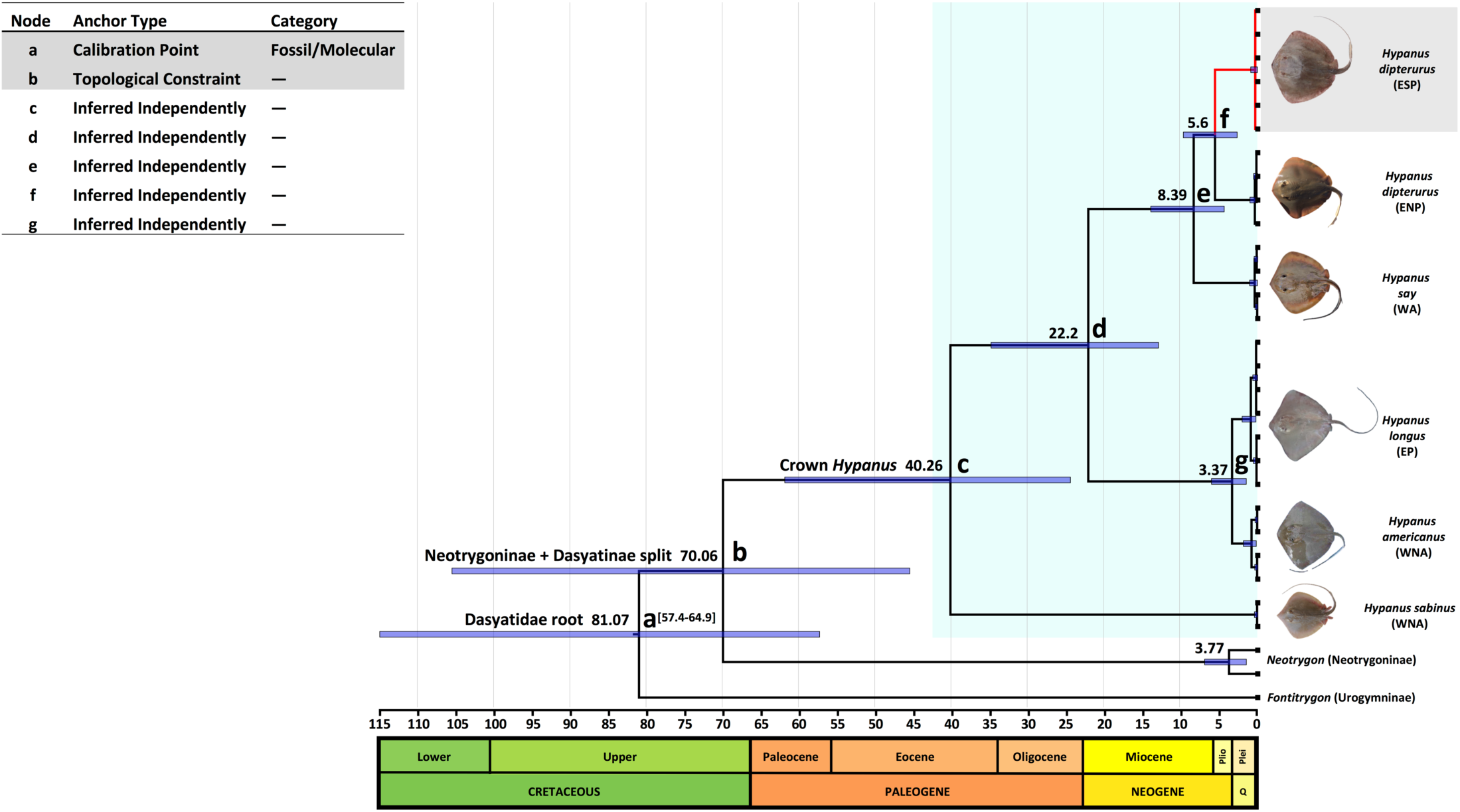
Time-calibrated Bayesian phylogeny illustrating the evolutionary relationships and estimated divergence of six target *Hypanus* species: *H. americanus*, *H. dipterurus*, *H. longus*, *H. sabinus*, and *H. say.* A concatenated dataset of 1,116 bp of the mitochondrial COI and 16S rRNA was used in the analysis. Node a: the root of the family Dasyatidae calibrated with an offsetlognormal distribution (57.4-64.9 Ma, indicated between brackets) based on Aschliman et al. (2012). Node b: hard topological constrain enforced to maintain the monophyly of Neotrygoninae and Dasyatidae clades following the consensus phylogeny of the family. Node c: crown *Hypanus* ingroup (subfamily Dasyatinae), shaded in light green color. Nodes d to g: divergence nodes inferred independently by the molecular clock. Outgroups include two *Neotrygon* species: *N. kuhlii* and *N. indica* (subfamily Neotrygoninae) and *Fontitrygon margarita* (subfamily Urogymninae). The ESP *H. dipterurus* clade, which is identified as a distinct linage that was split from the ENP *H. dipterurus*, is highlighted in red. Numbers at nodes represent the mean estimate divergence times in millions of years. Blue horizontal bars indicate the 95 % highest posterior density (HPD) intervals. The geological time scale at the bottom indicates the major eras, with periods ranging from the Cretaceous to the Quaternary (Q). Plei: Pleistocene Epoch, Plio: Pliocene Epoch. Species distributions: EP (Eastern Pacific), ESP (Eastern South Pacific), ENP (Eastern North Pacific), WA (Western Atlantic), WNA (Western North Atlantic). Photograph credits: image of ESP *H. dipterurus* by F. Zavalaga; image of *H. longus* was taken by L. Bancayán; images of *H. americanus*, *H. sabinus*, and *H. say* were kindly provided by R. Robertson (Robertson and Van Tassell 2023); image of ENP *H. dipterurus* was retrieved from the iNaturalist database (2026).

The radiation of the major *Hypanus* lineages, including the crown *Hypanus* (node c) and the ancestor of the clade containing *H. americanus*, *H. dipterurus*, *H. longus*, and *H. say* (node d) were independently inferred at 40.26 Ma (95% HPD: 24.54 – 61.96 Ma) and 22.2 Ma (95% HPD: 13.01 – 34.94 Ma), respectively. Independent inferences for transisthmian geminate pairs showed that the subclade *H. say*/*H. dipterurus* (node e) split at 8.39 Ma (95% HPD: 4.4 - 14.01 Ma), while the split of *H. americanus*/*H. longus* (node g) was estimated at 3.37 Ma (95% HPD: 1.51 – 6.06 Ma). Notably, the intra-Pacific divergence between the ENP and ESP *H. dipterurus* populations (node f) was independently inferred at approximately 5.6 Ma (95% HPD: 2.7 – 9.75 Ma). These findings suggest a long-standing pre-isthmian isolation between the *H. dipterurus* populations that support the existence of two different evolutionary units rather than a single continuous Eastern Pacific. Interestingly, the intra-Pacific divergence between ESP and ENP *H. dipterurus* populations (mean COI distance = 4 %) predated the transisthmian split of *H. americanus* and *H. longus* (mean COI distance = 2.5 %). While the latter fully encompasses the final closure of the Isthmus of Panama, the deeper genetic divergence within both *H. dipterurus* populations (see Table S1) further reinforces an earlier isolation event in the Eastern Pacific.

Although the double-anchored model improved ancestral node accuracy at deeper levels, it overestimated the substitution rates for recent mitochondrial divergences in transisthmian geminate pairs (Fig. S1). Notably, the divergence between *H. dipterurus* populations (node f) and the transisthmian split of *H. americanus*/*H. longus* (node g) were artificially compressed to 3.08 Ma and 2.26 Ma, respectively. While the single-anchored model estimated the *H. americanus*/*H. longus* divergence at 3.37 Ma—showing a remarkable correlation with the final closure of the Isthmus of Panama (Thacker 2017)—the alternative double-anchored model reduced this age to 2.26 Ma. This placement within the Pleistocene postdates the major oceanographic shifts and progressive shoaling of the Central American Seaway (CAS) occurring during the Pliocene (O’Dea et al. 2016). Consequently, these findings reinforce the single-anchored model as the most accurate representation of the recent evolutionary history of *Hypanus* lineages in the Eastern Pacific.

Based on the findings of this research, which strongly support that *H. dipterurus* populations from the ENP and ESP represent isolated evolutionary units, we propose that the ESP lineage be formally recognized as *H. brevis*. This taxonomic assignment follows Garman’s (1880) original description based on specimens from Paita, northern Peru. The revalidation of *H. brevis* increases the number of recognized species to 12 and brings the total number of distinct *Hypanus* evolutionary units to 15 (Table 4), including the three molecular evolutionary units (*H.* aff. *americanus, H.* aff. *guttatus,* and *H.* aff. *say*) identified by Petean et al. (2024).

**Table 4.** Current accepted species and latest taxonomic changes in members of the genus *Hypanus*. Distribution: EP (Eastern Pacific), ESP (Eastern South Pacific), ENP (Eastern North Pacific), ETP (Eastern Tropical Pacific), WA (Western Atlantic), EA (Eastern Atlantic), ECA (Eastern Central Atlantic), WCA (Western Central Atlantic), WNA (Western North Atlantic), WSA (Western South Atlantic), Gulf of Mexico (GM), NB (Northern Brazil), SB (Southern Brazil), and CS (Caribbean Sea).

| Taxonomy | Common name | Description Year | Distribution | Original name | Latest taxonomic change | Reference | IUCN status |
| --- | --- | --- | --- | --- | --- | --- | --- |
| **Genus** |  |  |  |  |  |  |  |
| *Hypanus* Rafinesque, 1818 | Stingrays | 1818 | EP, WA, EA | *Hypanus* Rafinesque, 1818 | Resurrected from synonymy with *Dasyatis* | Last et al. (2016b) | 2 as CR (18.1 %)<br>1 as EN (9.1 %)<br>3 as VU (27.3 %)<br>3 as NT (27.3 %)<br>1 as LC (9.1 %)<br>1 as NE (9.1 %) |
| **Validated species by the present work** |  |  |  |  |  |  |  |
| *Hypanus brevis* (Garman, 1880) | Peruvian diamond stingray | 1880 | ESP | *Trygon brevis* Garman, 1880 | Resurrected from synonymy with *H. dipterurus* | This study | VU (needs reassessment) |
| **Validated species in the literature** |  |  |  |  |  |  |  |
| *Hypanus guttatus* (Bloch & Schneider, 1801) | Longnose stingray | 1801 | WCA to SB | *Raja guttata* Bloch & Schneider, 1801 | Moved from the genus *Dasyatis* (*D. guttata*) | Last et al. (2016b) | NT |
| *Hypanus say* (Lesueur, 1817) | Bluntnose stingray | 1817 | WA | *Raja say* Lesueur, 1817 | Moved from the genus *Dasyatis* (*D. say*) | Last et al. (2016b) | NT |
| *Hypanus rudis* (Günther, 1870) | Smalltooth stingray | 1870 | ECA | *Trygon rudis* Günther, 1870 | Moved from the genus *Dasyatis* (*D. rudis*) | Last et al. (2016b) | CR |
| *Hypanus dipterurus* (Jordan & Gilbert, 1880) | Diamond stingray | 1880 | ENP | *Dasybatis dipterura* Jordan & Gilbert, 1880 | Moved from the genus *Dasyatis* (*D. dipterura*) | Last et al. (2016b) | VU (needs reassessment) |
| *Hypanus longus* (Garman, 1880) | Longtail stingray | 1880 | ETP | *Trygon longa* Garman, 1880 | Moved from the genus *Dasyatis* (*D. longa*) | Last et al. (2016b) | VU |
| *Hypanus sabinus* (Lesueur, 1824) | Atlantic stingray | 1824 | WNA | *Trygon sabina* Lesueur, 1824 | Moved from the genus *Dasyatis* (*D. sabina*) | Last et al. (2016b) | LC |
| *Hypanus americanus* (Hildebrand & Schroeder, 1928) | Southern stingray | 1928 | WNA, WCA and CS | *Dasyatis americana* Hildebrand & Schroeder, 1928 | Moved from the genus *Dasyatis* (*D. americana*) | Last et al. (2016b) | NT |
| *Hypanus geijskesi*<br>(Boeseman, 1948) | Wingfin<br>stingray | 1948 | WCA, NB | *Dasyatis*<br>*geijskesi*<br>Boeseman,<br>1948 | Moved from<br>the genus<br>*Fontitrygon* | Petean et al.<br>(2024) | CR |
| *Hypanus marianae*<br>(Gomes, Rosa &<br>Gadig, 2000) | Brazilian large-<br>eyed stingray | 2000 | WSA<br>(Brazil) | *Dasyatis*<br>*marianae*<br>Gomes, Rosa &<br>Gadig, 2000 | Moved from<br>the genus<br>*Dasyatis* (*D.*<br>*marianae*) | Last et al.<br>(2016b) | EN |
| *Hypanus berthallutzae*<br>Petean, Naylor &<br>Lima, 2020 | Lutz's stingray | 2020 | WSA<br>(Brazil) | *Hypanus*<br>*berthallutzae*<br>Petean, Naylor<br>& Lima, 2020 | New<br>discovery. It<br>was split<br>from *H.*<br>*americanus* | Petean et al.<br>(2020) | VU |
| *Hypanus rubioi*<br>Mejia-Falla, Navia,<br>Cardeñosa & Tavera,<br>2025 | Pointed-nose<br>Pacific stingray | 2025 | ETP | *Hypanus rubioi*<br>Mejia-Falla,<br>Navia,<br>Cardeñosa &<br>Tavera, 2025 | New<br>discovery | Mejia-Falla<br>et al. (2025) | NE |
| **Molecular evolutionary units (aff.)** |  |  |  |  |  |  |  |
| *Hypanus* aff.<br>*americanus* | – | – | WNA,<br>WCA, CS | – | – | Petean et al.<br>(2024) | – |
| *Hypanus* aff. *guttatus* | – | – | WCA | – | – | Petean et al.<br>(2024) | – |
| *Hypanus* aff. *say* | – | – | GM | – | – | Petean et al.<br>(2024) | – |

**Table 5.** Morphometric and meristic characteristics of *Hypanus brevis* (Garman, 1880) based on specimens collected in the southern Pacific Ocean off Peru, expressed in millimeters and as percentages of disc width (%DW). Data for the synt specimens MCZ S-422 from Paita, Peru. Additional comparisons are made with data from Garman (1913); Lamilla et al. (1995); and from the syntypes of *Hypanus dipterurus* Jordan and Gilbert (1880), from Beebe and Tee-Van (1941), an * Data corrected from the original publication. (–) Data not reported.

|  |  |  |  |  |  |  |  |  |  |  |  |  |  |  |
| --- | --- | --- | --- | --- | --- | --- | --- | --- | --- | --- | --- | --- | --- | --- |
| Sex | female | male | female | male | - | female | male | male | female | female | female | male | female | Male |
| Maturity | adult | juv | juv | juv | - | ? | young |  |  |  |  |  |  |  |

### Resurrection of *Hypanus brevis* (Garman, 1880)

Fig. 5 and Fig. 6; Table 5.

**Fig. 5.**
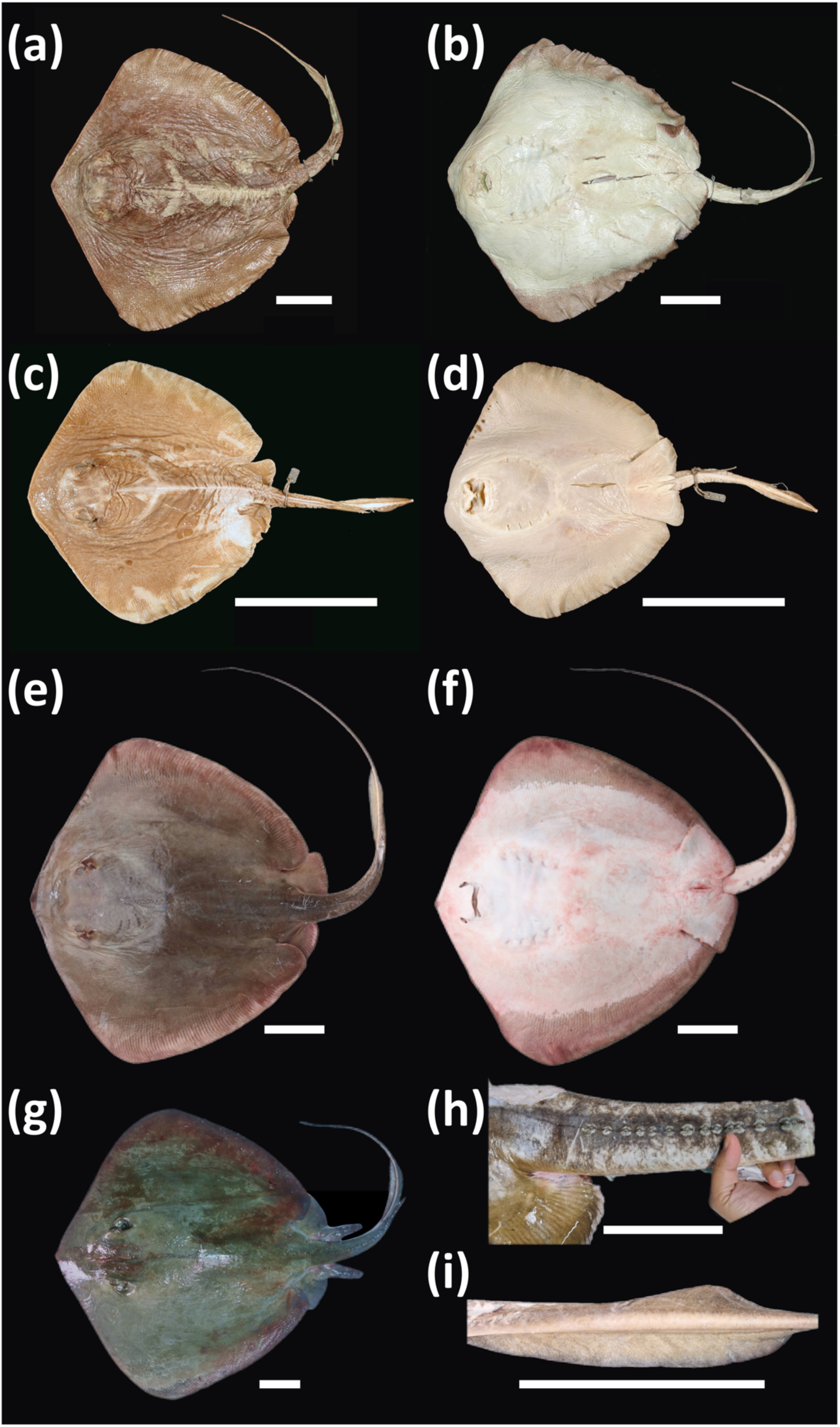
*Hypanus brevis* (Garman, 1880) from Peruvian waters. (a)–(d) Dorsal and ventral views of the syntypes of *Trygon brevis* Garman, 1880, female and male collected in Paita, MCZ S-422. (e)–(f) Dorsal and ventral views of female specimen IMARPE-20479 (present study). (g) Dorsal view of a male specimen collected in Chorrillos, MNHN-ICOS-02627 (TL = 1265 mm, DW = 690 mm). (h) Tail of specimen SV-0063 showing prominent, broad-based denticles (ca 900 mm DW). (i) Dorsal and ventral tail folds of specimen IMARPE-20479. Photographic credits: (a), (b), (c), and (d) Museum of Comparative Zoology, Harvard University; (e), (f), and (i) A. Moreno; (g) P. Béarez; and (h) F. Zavalaga. Scale bar: 10 cm.

**Fig. 6.**
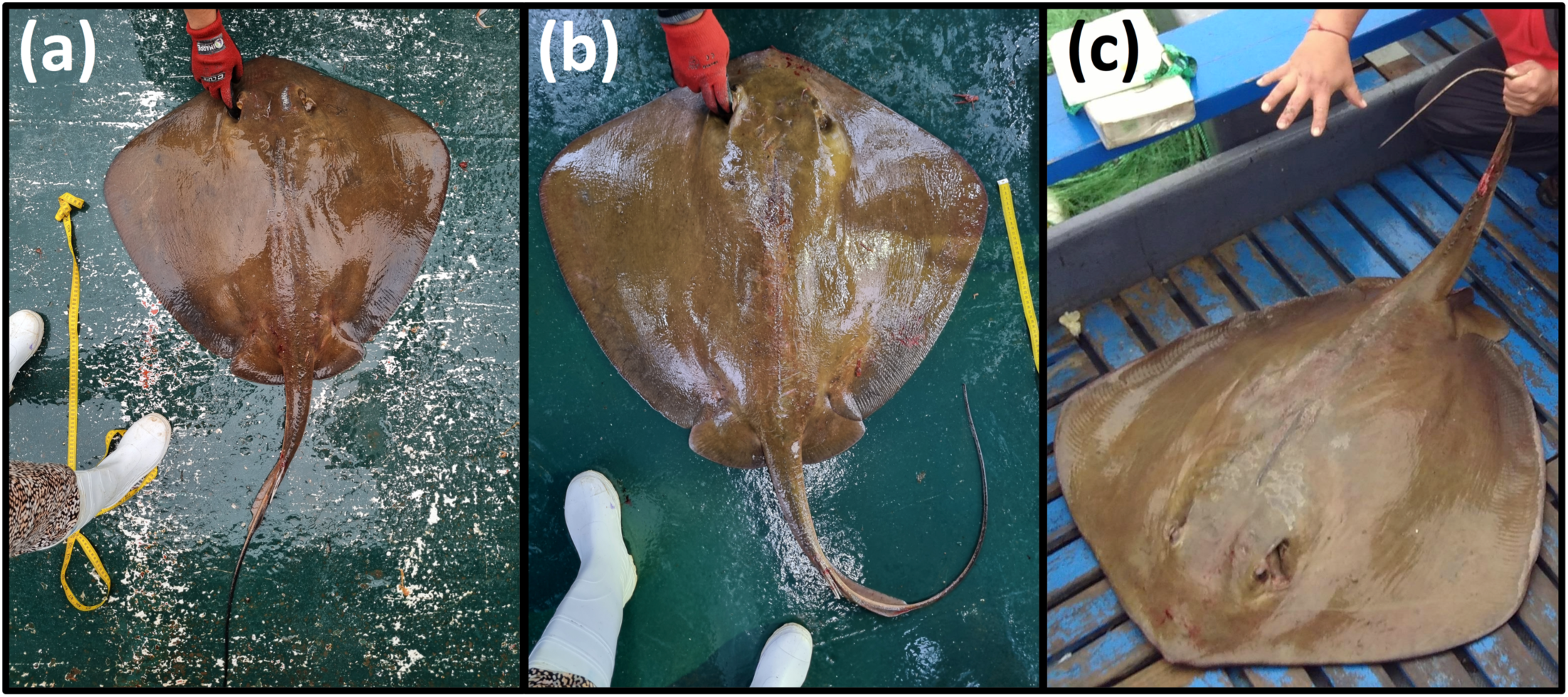
*Hypanus brevis* (Garman, 1880) Pictures illustrating the coloration observed in recently caught/live specimens. Panels (a) and (b): female specimens (SV-0146 and SV-0151, respectively) caught in San José (Lambayeque region, Peru); (c): specimen collected in Pisco (Ica region, Peru). Photographic credits: (a) and (b) F. Zavalaga; (c) J. Quiñones (used with written permission).

The reinstatement of *Hypanus brevis* is a nomenclatural act registered in ZooBank under the publication LSID: urn:lsid:zoobank.org:pub:7BAD4874-11C6-458C-AB18-0590D64A28B6.

*Trygon brevis* Garman, 1880: 171. Description.

*Dasibatis brevis* Garman (1883: 70).

*Dasybatus brevis*. In part Garman (1913: 396). Specimens of Paita, Peru, pl. 32, figs. 5-6.

*Dasyatis brevis* Chirichigno (1974: 75, figs. 67-68); Lamilla et al. (1995: 23-27);

Chirichigno and Vélez (1998: 65, figs. 11-12); Chirichigno and Cornejo (2001: 67).

*Hypanus dipterurus*. Almendras et al. (2025), specimens of Peru and Chile.

### Syntypes

MCZ Ichthyology S-422; 5°09′S, 81°08′W or −5.1583333°, −81.141666°; Paita, Piura, Peru; 5 May 1872; collected during the Hassler Expedition; held at Museum of Comparative Zoology, Harvard University. Fig. 5 (a)–(d).

Although Garman (1880) did not explicitly mention collection numbers, the existence of at least two syntypes is inferred from his description. He referred to a female specimen from Paita, Peru, and later confirmed that the two measured specimens (one male and one female) originated from Peru (Garman, 1883). No reference was made to type material from California.

### Diagnosis

A large-sized diamond-shaped species of *Hypanus* (attaining at least 900 mm DW) with the following combination of characters: body slightly wider than long; snout pointed but not elongated, rostral length 18.5–20.7 % DW; anterolateral margins of disc nearly straight, forming an angle of 124–130°; disc length 82.5–87.0 % DW; internostril width

8.5–9.7 % DW; oral papillae five, arranged in a transverse row on the floor of the mouth, with three well-developed medial papillae and one minute lateral papilla on each side; tail whiplike, less than 1.25 times disc width (1.4–1.5 times disc length), bearing a long, slender caudal spine and a short ventral caudal fin fold (25.7–34.6 % DW); dorsal disc generally smooth in juveniles, adults with scattered scapular denticles and enlarged denticles anterior to the caudal spine; coloration brown to olive-brown dorsally, lacking spots.

### Description

Disc wider than long, with disc width ranging from 241 to 540 mm and disc length from 206 to 470 mm, representing 82.55–87.04 % of disc width (DW). Anterolateral margins of disc nearly straight, posterior margins convex. Snout pointed but not protruding, with preorbital length ranging from 50 to 100 mm (18.52–20.75 % DW) and preoral length from 41 to 87 mm (15.77–17.01 % DW), forming an angle of 124–130°. Upper and lower jaws with small, blunt teeth. Oral papillae five, arranged in a single transverse row on the floor of the mouth; three medial papillae large and well developed, and one-minute lateral papilla on each side, widely separated from the medial group.

Pelvic fins broad, with pelvic fin length ranging from 42 to 104 mm (17.43–19.26 % DW) and width at base from 33 to 112 mm (12.75–20.74 % DW); margins truncate, apices slightly rounded, and angles rounded. Tail moderately long, measuring 289–663 mm, representing 119.92–123.99 % DW and 140.29–150.18 % of disc length (DL); broad and slightly depressed at the base, progressively tapering toward the caudal spine. Tail width at the spine base ranges from 6 to 16 mm (2.18–2.96 % DW).

Dorsal and ventral caudal folds present; dorsal fold low and short, measuring 34–78 mm in length (10.59–15.77 % DW) and 3–10 mm in height (1.17–1.85 % DW). Ventral fold higher and longer than dorsal, ranging from 62 to 187 mm in length (25.73–34.63 % DW) and from 3.5 to 16 mm in height (1.34–2.96 % DW) (Table 5 and Fig. 5). The ventral fold originates at the level of the caudal spine base, whereas the dorsal fold originates at the level of the caudal spine tip. Both folds terminate at approximately the same level along the tail, from where the tail becomes laterally compressed and gradually narrows into a filament (see Fig. 5i).

Dorsal surface of disc smooth. The largest specimen (IMARPE-20479), a female of 540 mm DW, exhibits an irregular row of scattered denticles extending from the nuchal region to the origin of the caudal spine, as well as a short row on each shoulder, with up to five denticles per side (Fig. 5e-f). Juvenile specimens lack denticles. Additionally, a large incomplete specimen collected at ∼6° S (SV-0063) showed more than 14 prominent, broad-based denticles on the tail anterior to the caudal spine (see Fig. 5h).

### Comparison

*Hypanus brevis* is distinguished from *H. dipterurus* by presenting a disc clearly wider than long, with disc length representing 82.5–87.0 % DW (vs. 88.9 to 94.9 % DW) and by a notably smaller internostril width, 8.5–9.7 % DW (vs. 13.6–18.6 % DW). However, both species overlap in several external characters, including a very short tail, measuring 1.4–1.5 times disc length (vs. 1.1–1.4 times); a small ventral caudal fin fold, 25.7–34.6 % DW (vs. 28.4–33.3 % DW); and a short, non-elongated rostrum, with rostral length of 18.5–20.7 % DW (vs. 15.9–21.8 % DW), as well as a wide rostral angle.

*Hypanus brevis* is readily distinguished from *H. longus* and *H. rubioi* by its much shorter tail, 1.4–1.5 times disc length (vs. 2.4 and from 2.4 to 3.1 times, respectively). Also, it shares with *H. longus* a disc wider than long, 82.5–87.0 % DW and 81.6–86.8 % DW, respectively. In contrast, *H. rubioi* presents a proportionally longer disc (89.6 to 98.2 % DW), nearly as long as wide.

The rostrum of *H. brevis* is short and not pointed, differing markedly from the elongated and pointed rostrum of *H. rubioi* with 29.5–32.1 % DW of rostral length. In contrast, *H. longus* shows rostral length values (16.4–19.5 % DW) similar to those of *H. brevis*. However, the rostral angle is slightly narrower in *H. longus*, whereas it is acute in *H. rubioi*. Likewise, *H. longus* and *H. rubioi* differ from *H. brevis* in having a much longer ventral caudal fin fold, measuring 45–52.4 % DW and 38–60.8 % DW, respectively. Internostril width in *H. longus* (9.4–10.4 % DW) and *H. rubioi* (9.1–10.7 % DW) is similar to, though slightly greater than, that of *H. brevis*.

### Color in life

Dorsal surface uniformly brown, dark brown, or olive-brown, becoming slightly reddish near the margins (see Fig. 6); ventral surface white, with the posterior margins of the pectoral and pelvic fins brown. Juveniles exhibit a slightly more violaceous coloration on the dorsal surface.

### Size

The largest known specimen of *H. brevis* is an incomplete mature individual of approximately 900 mm DW (SV-0063, present study), collected off Peru at ∼6°S (Fig. 5h). A male specimen 690 mm DW collected in Chorrillos, Lima, Peru, on 25 March 1999 (MNHN-ICOS-02627; Fig. 5g) weighed 13800 g. In the present study, the specimens examined measured between 241 and 540 mm DW.

### Distribution

Recent evidence suggests that *H. brevis* is restricted to the ESP, including commercial fisheries off Ecuador (Martínez-Ortiz and García-Domínguez 2013; Calle-Morán and Béarez 2020) and Colombia (Mejía-Falla and Navia 2019). These records may correspond to the northernmost occurrence of *H. brevis*; however, further genetic analyses are required to determine whether the diamond stingrays captured in these countries belong to *H. brevis* or *H. dipterurus*. To date, *H. brevis* has been genetically confirmed only from specimens collected in Peru (Marín et al. 2018; present study). Southward, several studies have reported the presence of *H. brevis* south to Antofagasta, northern Chile (Lamilla et al. 1995; Almendras et al. 2025), which likely represents the southernmost limit of its distribution.

### Common name

To differentiate *H. brevis* from its sister species, *H. dipterurus* (commonly known as the Diamond stingray), we propose the common name “Peruvian diamond stingray”. This name is fitting because the original description is based on specimens collected from northern Peru. As for the Spanish common name, the vernacular term “*batea*” is widely used in Peru as the commercial name for this particular species, and we suggest keeping it unchanged.

### Fishery

Over 29 years, a total of 135 thousand tons of chondrichthyans were recorded in the IMARPE landing database. These figures account for 19 % of the batoid group, with less than 1 % potentially attributable to *H. brevis* (top panel of Fig. 7). Notably, 98 % of these volumes were originally labeled by IMARPE as *Dasyatis brevis*, while the remaining were identified as *Hypanus* sp. Between 2015 and 2024, IMARPE recorded an annual catch of approximately 46.2 tons of *H. brevis* (*sd* of 12.3 tons), corresponding to an annual economic value of approximately 62,000 USD (*sd* of ± 16,000; see Fig. S2). The majority of the landings were reported from northern regions of the country (44 %; averaging 20.4 ± 6.6 tons per year) and from the southern regions (37 %; averaging 16.8 ± 9.8 tons per year; bottom panel of Fig. 7). During this period, 78 % of landings originated from gillnets, with the remaining percentage from unspecified fishing gear types other than gillnets (e.g., beach seines and purse seines as reported by González-Pestana et al. 2022).

**Fig. 7.**
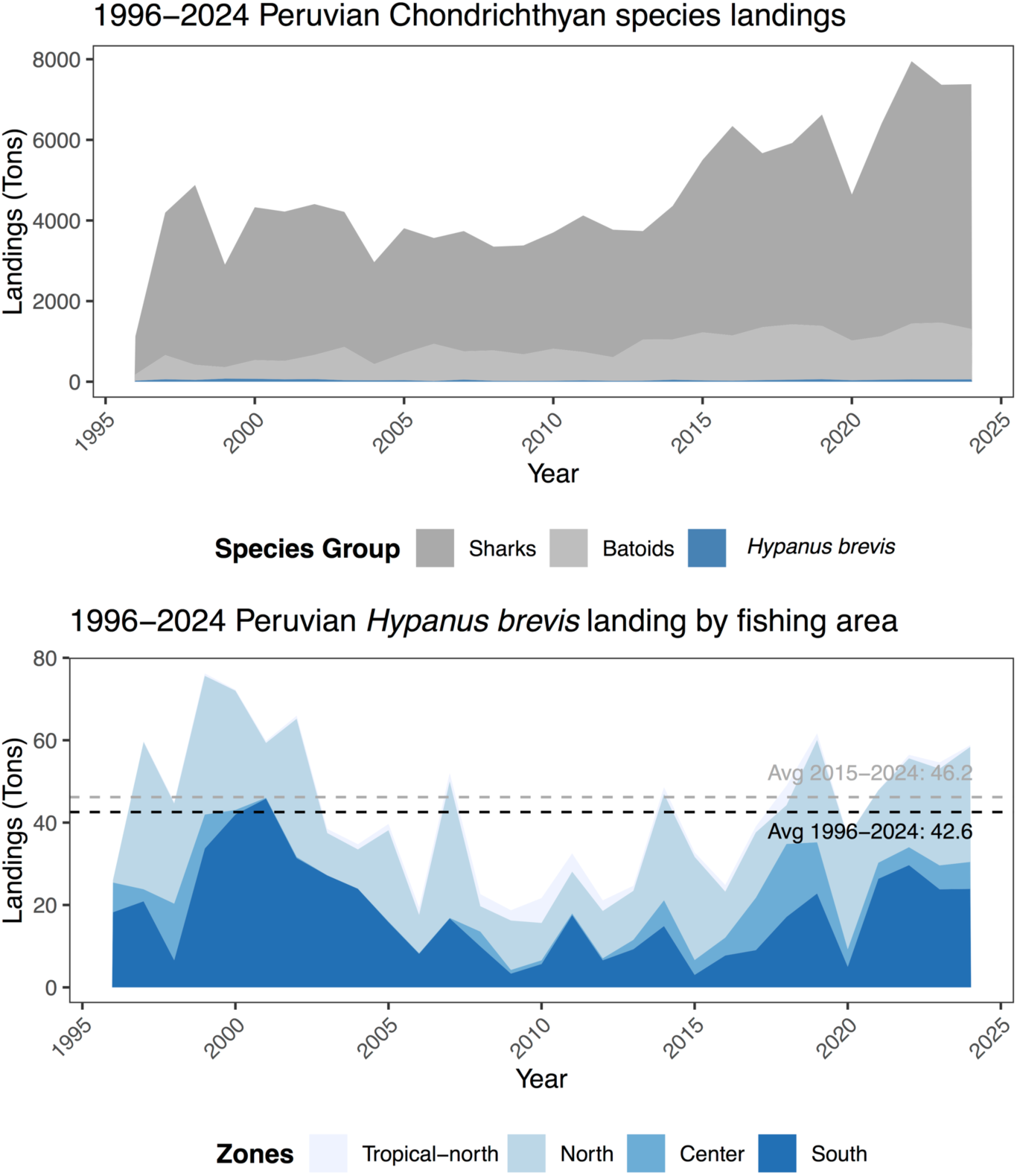
Peruvian Chondrichthyan species documented in the IMARPE landing data from 1996 to 2024 (top panel) and landings potentially attributable to *Hypanus brevis* within the same period (bottom panel).

The fishery for *H. brevis* is primarily coastal, with 32 % of reported catches occurring during 2015-2024 within the first nautical mile (nm) off the Peruvian coastline and an additional 44 % between 1-5 nm. The remaining catches, reported far from the 5 nm, are likely still within the continental shelf, as most of these catches occur around the *Lobos de Afuera* Islands (Fig. 8).

**Fig. 8.**
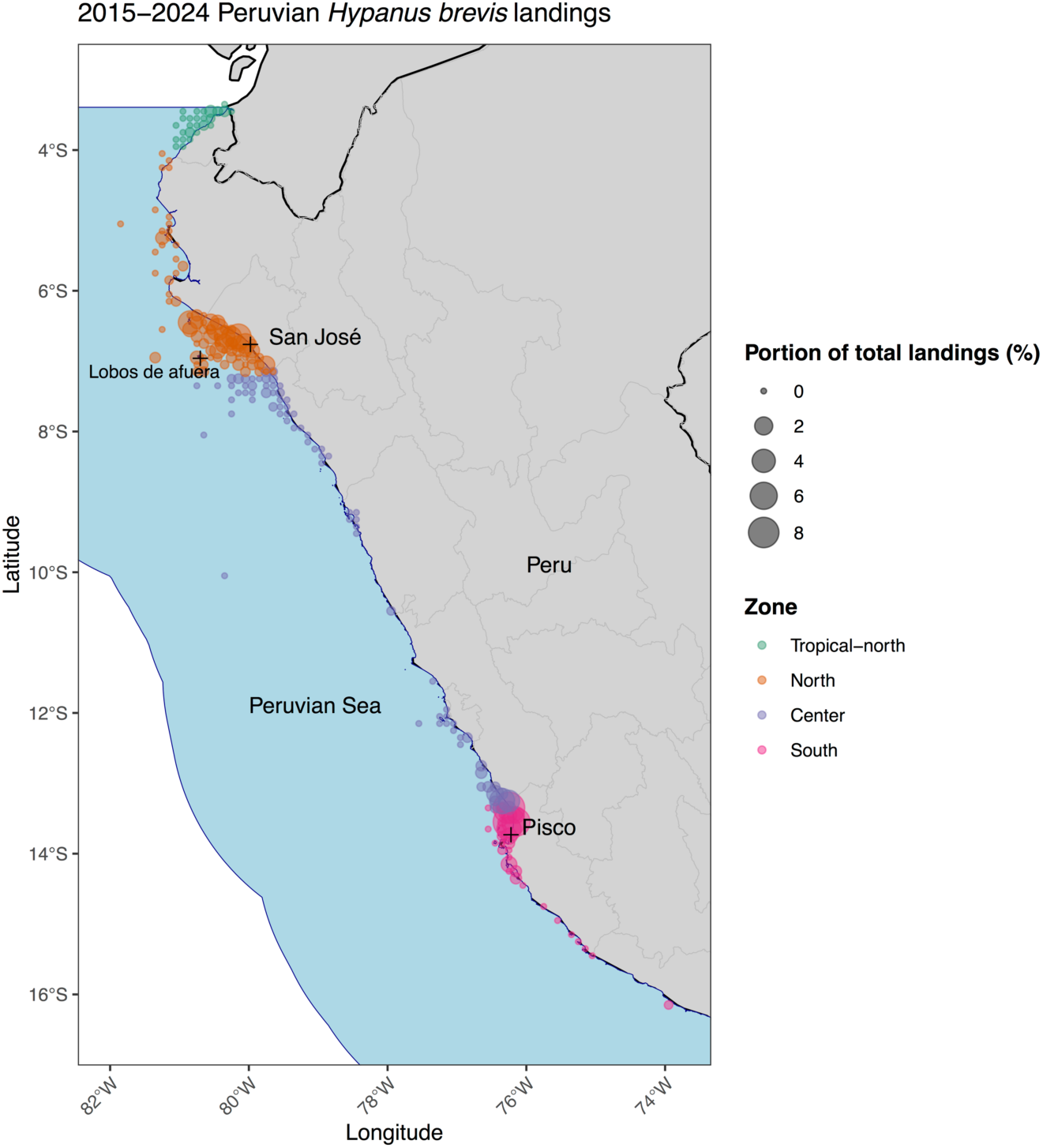
The occurrence of landings of *Hypanus brevis* in the Peruvian Sea from 2015 to 2024, classified within a 0.1° latitude by 0.1° longitude grid.

## 4. DISCUSSION

Allopatric speciation is a major evolutionary model that has shaped the population history of most aquatic species, including elasmobranchs (Ball et al. 2016; Gayford and Jambura 2025). This evolutionary mechanism has been a primary driver of species richness in batoids (Staggi et al. 2025), which comprises more than a half of all elasmobranch diversity (Ebert et al. 2017). The speciose nature of this elasmobranch group, coupled with its highly conserved morphology (dorsoventrally flattened bodies and countershaded coloration), have led to a long history of inaccurate taxonomic descriptions (Last et al. 2016b; Chatzispyrou and Koutsikopoulos 2025).

This issue is particularly evident in the genus *Hypanus*, whose taxonomic composition has been subjected to major controversy, with significant revisions performed since the last decade (Last et al. 2016b). Most of these taxonomic revisions (see Table 4) were accomplished by means of integrative taxonomic approaches, including DNA information from complete mitogenomes or partial gene sequences (Last et al. 2016b; Petean et al. 2020, 2024; Mejía-Falla et al. 2025). Nevertheless, the species diversity of *Hypanus* is far from taxonomic clarity, primarily due to unresolved species complexes and hidden lineages, warranting additional revisions (Petean et al. 2024).

The findings of the present research resolve a taxonomic debate spanning nearly one and a half century, originating with contemporaneous descriptions published by Jordan and Gilbert (1880) and Garman (1880). Through an integrative taxonomic approach combining robust molecular data (Table 3, Fig 2 to Fig. 4) with subtle but consistent morphological trends (Table 5), this study resurrects the Peruvian diamond stingray, *H. brevis* (ESP), from synonymy with *H. dipterurus* (ENP). The resurrection of *H. brevis* as a distinct evolutionary unit from the ESP was supported by the results of different molecular analyses conducted herein.

The haplotype network analyses (Fig. 2) revealed a complete lineage sorting, with no haplotypes shared between these two sister species. The Bayesian phylogenetic analysis (Fig. 3) recovered all specimens of *H. brevis* in a distinct monophyletic clade. The mean COI distance of 4 % observed between *H. dipterurus* and *H. brevis* (Table S1) is notably higher than the lowest average interspecific threshold (0.59 %) reported for congeneric taxa *H. berthalutzae* and *H. rudis* (Petean et al. 2024), reinforcing its validation as different species. The mean COI distance observed between *H. dipterurus* and *H. brevis* aligns with the inferred molecular divergence age estimated at 5.6 Ma (Fig. 4). The species status of both Pacific diamond stingrays was further supported by AMOVA results (Table 3). The high and significant Φ*_ST_* values confirm that the vast majority of genetic variation is partitioned between groups (ENP vs. ESP), indicating complete genetic isolation and lack of gene flow between them.

Morphologically, *H. brevis* and *H. dipterurus* appear difficult to distinguish. Both species share a generally similar disc shape, proportional measurements, dentition patterns, and external coloration, resulting in considerable overlap in the characters traditionally used for identification. Subtle differences have been suggested in previous studies; however, these traits often show intraspecific variation and may be influenced by ontogeny or preservation condition, reducing their diagnostic reliability. Consequently, the identification of these species based solely on external morphology remains challenging and may lead to misidentifications in the absence of comprehensive comparative material or additional lines of evidence.

Garman (1880, 1883) and Lamilla et al. (1995) reported the presence of five oral papillae in *H. brevis*. The present study likewise confirms five papillae on the floor of the mouth, arranged in a transverse row, consisting of three well-developed medial papillae and one-minute lateral papilla on each side, widely separated from the medial group. The small size and inconspicuous nature of the lateral papillae may hinder their proper recognition. Therefore, although the number of papillae appears consistent, careful examination is required when assessing this character.

Dermal ornamentation in *H. brevis* appears to be strongly influenced by ontogeny and body size rather than representing a stable diagnostic character. Garman (1880, 1883), based on a large female specimen, described three large, erect, broad-based denticles anterior to the caudal spine, together with a roughened tail bearing smaller tubercles. A comparable condition was observed in the present study in a large incomplete specimen (SV-0063), approximately 900 mm DW, which exhibited more than 14 well-developed denticles on the tail anterior to the caudal spine (Fig. 5h), whereas smaller juvenile specimens lacked denticles entirely.

Short rows of one to four tubercles on the shoulder region were also noted by Garman (1880, 1883), suggesting that dermal denticles increase in number and prominence with growth, as reported for related Atlantic congeners. A similar pattern was reported by Beebe and Tee-Van (1941) in *H. dipterurus* from Mexico, where larger individuals exhibited more conspicuous dorsal denticles and scapular tubercles. This suggests that dermal ornamentation may vary with ontogeny and possibly sex, although this character remains poorly documented in both species. Therefore, dermal ornamentation should be interpreted cautiously and not considered a primary diagnostic character for separating *H. brevis* from *H. dipterurus*, and its taxonomic significance requires further study.

With respect to coloration, early descriptions already suggested differences among *Hypanus* specimens along a latitudinal gradient. Jordan and Gilbert (1880) described *H. dipterurus* specimens as light brown, slightly marbled with darker pigmentation, lacking distinct spots, and with a blackish tail. Furthermore, Beebe and Tee-Van (1941) reported that specimens from the ENP exhibited a darker overall coloration, ranging from dark brown to black. In contrast, the specimens of *H. brevis* examined in the present study from Peru showed a generally brown to olive dorsal coloration, sometimes with slightly reddish tones, and a distinctly paler ventral surface.

Although these differences in pigmentation may reflect geographic variation and could support separation between *H. brevis* and northern *H. dipterurus*, coloration alone should be interpreted cautiously, as it may vary with preservation, ontogeny, and environmental conditions. Therefore, pigmentation is better considered a complementary rather than a primary diagnostic character, and broader comparative studies across the distributional range of both species are needed to assess its taxonomic significance.

Regarding body size, specimen SV-0063 represents the largest documented record of *H. brevis* directly observed by the authors, reaching approximately 900 mm disc width (DW) (Fig. 5h). Chirichigno and Cornejo (2001) reported maximum total lengths of up to 1.8 m TL. Although these larger sizes were not verified through examined material, they suggest that *H. brevis* may attain greater dimensions than currently represented in scientific collections.

The scarcity of large specimens in reference collections likely reflects logistical difficulties in capturing, handling, and preserving large rays rather than true rarity in nature. As a result, *H. brevis* remains poorly represented in museum collections and historically understudied. Similar limitations have been reported for other Eastern Pacific chondrichthyans, where insufficient material constrains robust taxonomic and biological assessments (Becerril-García et al. 2022).

It has been proposed that the history of the Pacific *Hypanus* lineages started during the Pliocene, with the formation of the Isthmus of Panama (Rosenberger 2001). This major geological event occurred about three to four Ma (Thacker 2017) and may have led to the allopatric speciation of three Pacific *Hypanus* lineages: *H. dipterurus*, *H. longus*, and *H. rubioi*, which branched off from their Atlantic transisthmian geminates: *H. say*, *H. americanus*, and *H. guttatus*, respectively (Rosenberger 2001; Petean et al. 2024; Mejía-Falla et al. 2025). To get further insights into their evolutionary history, we employed the combined information from the COI and 16S rRNA genes to estimate their divergence times. Regrettably, the lack of 16S rRNA sequences in public databases for the recently discovered *H. rubioi* prevented us from dating its transisthmian divergence.

Our molecular dating analysis, based on the primary single-anchored model (Fig. 4), revealed a distinct and independent evolutionary history among the transisthmian *Hypanus* lineages. While the split between *H. longus* and *H. americanus* was estimated at 3.37 Ma—coinciding with the final stages of the CAS closure—the divergence of *H. dipterurus* from *H. say* was inferred at 8.39 Ma. These results support a “pre-isthmian divergence” hypothesis for the latter geminate pair, suggesting that the CAS closure did not affect all *Hypanus* lineages simultaneously. Instead, vicariance was likely staggered, driven by differential habitat preferences and ecological traits. The inferred split between *H. dipterurus* (ENP) and *H. brevis* (ESP) lineages was estimated at approximately 5.6 Ma, suggesting that the gene flow was interrupted during the late Miocene/early Pliocene. These findings support the hypothesis that the monophyletic amphi-American lineage comprised by *H. brevis*, *H. dipterurus*, and *H. say*, emerged through pre-isthmian vicariance (Knowlton and Weigt 1998), followed by a secondary antitropical intra-Pacific radiation (Thacker 2017). This two-step speciation model can be attributed to geological and oceanographic shifts that occurred during the gradual shoaling of the CAS, which lasted for approximately 12 million years (Lessios 2008).

The *H. brevis*/*H. dipterurus* divergence, estimated at 5.6 Ma, aligns with the Late Miocene Biogenic Bloom, a period characterized by the intensification of Pacific upwellings, the influx of cooler, nutrient-rich deep waters, and salinity divergence (O’Dea et al. 2016; Carrillo-Briceño et al. 2018; Reghellin et al. 2022). This oceanographic reconfiguration during the early stages of the CAS shoaling established initial thermal barriers, interrupting gene flow in several marine taxa (Lessios 2008). This early isolation was subsequently reinforced and finalized by the climatic oscillations of the Pliocene and Pleistocene. During the glacial periods, cold-water zones acted as “thermal corridors”, enabling marine taxa to cross the Equator, while subsequent warm interglacial waters functioned as “thermal barriers” (Burridge 2002; Ludt 2021). These cyclic shifts pushed ancestral Pacific populations toward the poles, effectively leading to allopatric speciation (Lindberg 1991).

The proposed “tropical gap” between *H. brevis* and *H. dipterurus* is currently based on a lack of confirmed occurrences in Central America. Our study analyzed genetically verified samples from the core populations of both species: *H. brevis* from the ESP (Peru) and *H. dipterurus* from the ENP (Mexico and southern USA). These locations represent typical temperate to subtropical waters, as defined by established marine ecoregions (Spalding et al. 2007). Although previous studies have reported low catch frequencies of *H. dipterurus* in Central America (e.g., Panama, Morales-Saldaña et al. 2025), these records may represent misidentifications. It is also likely that infrequent sightings of *H. dipterurus* in Central American waters are merely transient range shifts triggered by climatic phenomena (Cerutti-Pereyra et al. 2024). In this light, further research is needed to confirm the actual geographic distribution boundaries of these stingrays.

Notably, a total homogeneity in the COI diversity was observed in all analyzed *H. brevis* samples (n =27). A founder event could not only be a possible speciation pathway in *H. brevis* but also it would explain the low intraspecific diversity in *H. brevis*, where a single, fixed haplotype was detected in the COI gene. However, the divergence span of 5.6 Ma, suggested by our molecular dating analysis, is more than enough for new mutations to accumulate and establish different haplotypes (Martin et al. 1992; Dos Reis et al. 2015). The fixation of a single COI haplotype across a wide latitudinal gradient (∼3°S to 14°S), from the tropical waters of Tumbes to the temperate-cold waters of Pisco, suggests high connectivity among the Peruvian samples of *H. brevis*. High levels of gene flow among these individuals are likely facilitated by the northward movement of the Humboldt Current System (HCS) (Thiel et al. 2007).

The reduced genetic diversity of *H. brevis* is of particular conservation concern given that low genetic variation is often associated with a reduced adaptive potential to climatic changes (Reed and Frankman 2003). While our genetic results indicate that *H. dipterurus* from the ENP showed classic signs of population expansion, *H. brevis* from the ESP, contrastingly, may have experienced a recent population bottleneck. This warrants immediate research on the ESP populations of the recently resurrected Peruvian diamond stingray lineage. These studies should conduct population genetic structure analyses and the estimation of effective population size using high-resolution genomic approaches with additional nuclear markers in more intensive geographic sampling. The results from these studies will determine whether the observed low mitochondrial variation in *H. brevis* reflects a critical population bottleneck or widespread connectivity within the Peruvian populations. Ultimately, these data will clarify the population’s adaptive potential and enable the identification of management units needed for robust conservation strategies.

### Taxonomic justification for the resurrection

Our specimens agree closely with the original description of *Trygon brevis* Garman (1880) currently recognized as *Hypanus brevis*, and with the syntype material (MCZ Ichthyology S-422) in several diagnostic features, particularly the short tail, the disc clearly wider than long, the short and non-elongated snout, the broad rostral angle, and the relatively small ventral caudal fold (see Table 5). These characters clearly separate this lineage from *H. dipterurus*, with which *H. brevis* has historically been synonymized (see Comparison section).

Although the original description of *Trygon brevis* provides only limited morphometric information suitable for direct comparison, mainly disc width (DW), disc length (DL), and caudal length (CL), these measurements show broad agreement with the present material.

Garman (1880) reported disc length values of 88.06–94.44 % DW and caudal length values of 127.77–130.42 % DW for the syntypes, whereas our specimens showed disc length ranging from 82.55 to 87.04 % DW and caudal length from 119.92 to 123.99 % DW (mean = 84.99 and 121.87 % DW, respectively; Table 5). Although the syntypes present slightly higher values, both datasets consistently support the same general morphology: a disc distinctly wider than long and a relatively short tail, corresponding to Garman’s original diagnosis of a “tail less than one and a half times the length of the disk.” In addition, the qualitative description matches remarkably well with the present material. Garman (1880) described the species as having a “disk quadrangular, a little wider than long,” short and broad caudal folds, a blunt snout, rounded pectoral extremities, and broad ventral fins, all of which are consistent with our specimens (see Description section). The presence of enlarged scapular tubercles and caudal denticles in adults also agrees with the ontogenetic variation observed in the present material and in other species of *Hypanus* (Figure 5).

Likewise, Chirichigno (1974), Lamilla et al. (1995), Chirichigno and Vélez (1998), and Chirichigno and Cornejo (2001) recognized specimens from the southeastern Pacific matching this morphology and questioned the synonymy with *H. dipterurus*. Our morphometric comparisons support this interpretation, particularly in disc proportions and internostril width, which consistently distinguish the Peruvian lineage from *H. dipterurus*. Although some variation exists between our material and the original description, these differences are minor and are likely attributable to preservation effects, ontogenetic variation, and differences in measurement protocols between historical and modern studies. The scarcity of quantitative morphometric data in the original description limits a more detailed direct comparison; however, the strong concordance in diagnostic external morphology supports the interpretation that the lineage examined here corresponds to the original concept of *Hypanus brevis* rather than representing an undescribed species. Therefore, the resurrection of *Hypanus brevis* is herein supported.

### Conservation and management considerations

The taxonomic separation of *H. brevis* from *H. dipterurus* indicates that both species possess a more restricted distribution than previously understood when they were considered as a single species. Based on the available information, Peru serves as a significant hub for the fishing of *H. brevis* by small-scale operators, with certain reports indicating the presence of this species in northern Chile (Bustamante et al. 2023; Almendras et al. 2025) and Ecuador, including the Galápagos Islands (Grove et al. 2022; Victor et al. 2022; Bermello-Vera 2024; Alcivar-Holguin and Pilligua-Holguin 2025). Moreover, further research is required to validate the distribution range and population structure, and to ascertain the exploitation status of the stock(s). *Hypanus dipterurus* is a species that exhibits localized depletion due to its intrinsic life-history traits and excessive fishing pressure (Smith et al. 2008). Given their close relatedness, the latter underscores the need to protect the two main fishing spots in Peru (see Fig. 8), in a way to prevent future collapses of *H. brevis*, which exhibits remarkable mitochondrial homogeneity (see a discussion of area-based management mechanisms in Campos-León et al. 2025).

Considering the relative importance of the Peruvian fishery as the sole focus of regular fishing efforts throughout the years, it is a positive signal the fact that Peruvian landings have not experienced a sharp decline over the past three decades (Fig. 7). Nonetheless, this should not be interpreted in isolation as an indicator of sustainable fishing practices. While catches may have remained relatively stable and provided a relatively consistent supply to markets over time, the unit of fishing effort may have increased, and/or the average size of caught individuals may have decreased. IMARPE maintains a comprehensive monitoring program for artisanal fisheries that likely contains relevant data that could thereby facilitate answering some of the relevant research questions to inform specific management provisions. Therefore, it is advisable to disclose existing biological and fisheries data and to develop plans for the collection of supplementary data (González-Pestana et al. 2022; Campos-León et al. 2025). Revision of the available datasets may lead to an update of the life-history traits, as until now most have been estimated solely through sampling *H. dipterurus* individuals within the ENP (Mariano-Meléndez 1997; Smith 2005; Smith et al. 2008; Restrepo-Gómez et al. 2020), while fewer studies have been conducted on the ESP *H. brevis*. With some exceptions, the life-history parameters of *H. brevis* remain largely undocumented (Silva-Garay et al. 2018; Sánchez-Rea and Kanagusuku 2020; Garcia-Yarihuamán and Mantarí-Gavilano 2021; González-Pestana et al. 2022;).

Finally, *H. dipterurus* is classified as a vulnerable species in its most recent assessment by the IUCN Red List of Threatened Species, considering that “[i]n Peru, directed gillnet fisheries landings for this species have seen a decline of 50 % in annual landed biomass between 1997 and 2015 […]” (Pollom et al. 2020). Our results, covering a longer period than the IUCN evaluation, do not show a declining landing trend. However, this should not be construed as an indicator of sustainable management, given that the fishery is undermanaged. Therefore, the absence of a clear definition of the exploitation status should not justify failure to implement management measures; instead, a more precautionary approach must be adopted, acknowledging the significant uncertainty and recognizing that the genetic diversity of the reclassified *H. brevis* is impoverished, serving as a potential proxy for population health that indicates conservation concerns. Consequently, it calls for further analyses to identify management strategies that prevent additional losses of genetic variation. Future IUCN evaluations should consider *H. brevis* as an independent species.

## AUTHOR CONTRIBUTIONS

### Alan Marín

conceived and designed the study, supervised the project, designed the molecular experiments, analyzed the data, conducted the population genetics/phylogenetic/evolutionary analyses, developed figures, wrote the first draft of the manuscript.

### Fabiola Zavalaga

conceived and designed the study, conducted field work, conducted and led the morphometric analysis, analyzed the data, developed figures, wrote the original draft (morphometric sections).

### Renato Gozzer-Wuest

conceived and designed the study, conducted the fisheries analysis, analyzed the data, developed figures, wrote the original draft (fisheries and conservation sections).

### Luis E. Santos-Rojas

conducted field work, performed the molecular experiments, analyzed the data.

### Lorenzo E. Reyes-Flores

conducted field work, performed the molecular experiments, analyzed the data.

### Ruben Alfaro

conducted field work, performed the molecular experiments, analyzed the data.

### Philippe Béarez

conceived and designed the study, conducted and led the morphometric analysis, analyzed the data, developed figures, reviewed and edited the manuscript, wrote the original draft (morphometric sections).

### Eliana Zelada-Mázmela

conceived and designed the study, reviewed and edited the manuscript, provided laboratory facilities for molecular analysis, administered the project, responsible for funding acquisition.

All authors read, revised, and approved the final version of the manuscript.

## ACKNOWLEDGEMENTS

The authors thank Luis Bancayán for providing photographic material and tissue samples of *H. longus* from Tumbes, Peru. We are grateful to R. Robertson and J. Quiñones for providing photographs of *H. americanus*, *H. brevis*, *H. sabinus*, and *H. say*. Special thanks are also extended to fisheries biologist Andrey Moreno for his assistance with the photographic documentation of *H. brevis* specimens deposited at IMARPE, as well as for his support in specimen handling.

## DATA AVAILABILITY STATEMENT

All DNA sequences generated in the present study are available in the GenBank database under accession numbers from PZ027578 to PZ027634.

---

**Fig. S1.**
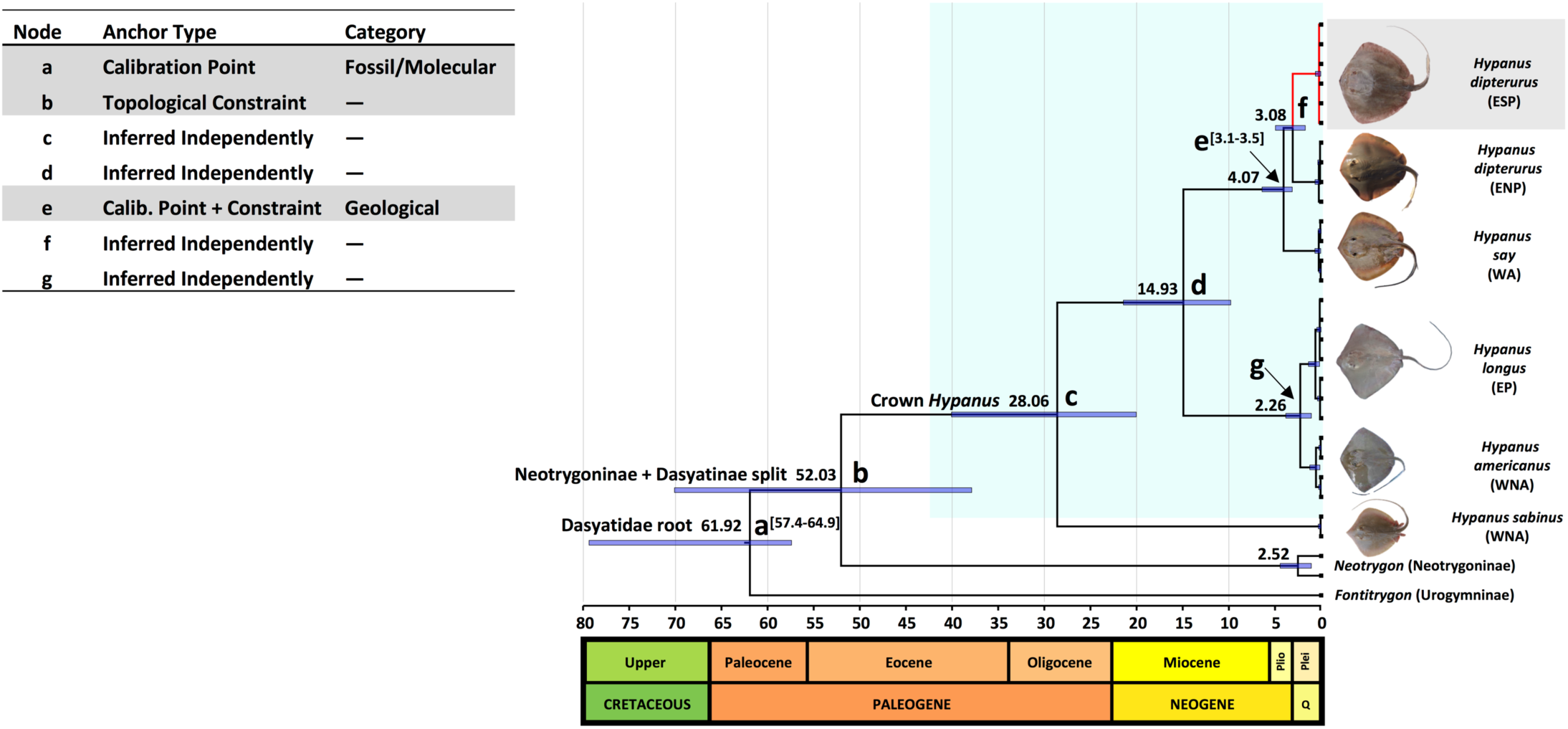
Time-calibrated Bayesian phylogeny illustrating the sensitivity analysis chronogram of the family Dasyatidae based on a concatenated dataset of 1,116 bp of the mitochondrial COI and 16S rRNA and double-anchored calibration approach. Node a: the root of the family Dasyatidae calibrated with an offsetlognormal distribution (57.4-64.9 Ma, indicated between brackets) based on Aschliman et al. (2012). Node b: hard topological constrain enforced to maintain the monophyly of Neotrygoninae and Dasyatidae clades following the consensus phylogeny of the family. Node c: crown *Hypanus* ingroup (subfamily Dasyatinae), shaded in light green color. Node e: internal geological calibration point representing the final closure of the Isthmus of Panama (3.1-3.5 Ma, indicated between brackets) based on O’Dea et al. (2016). Nodes c, d, f, and g were inferred independently. Outgroups include two *Neotrygon* species: *N. kuhlii* and *N. indica* (subfamily Neotrygoninae) and *Fontitrygon margarita* (subfamily Urogymninae). The ESP *H. dipterurus* clade, which is identified as a distinct linage that was split from the ENP *H. dipterurus*, is highlighted in red. Numbers at nodes represent the mean estimate divergence times in millions of years. Blue horizontal bars indicate the 95 % highest posterior density (HPD) intervals. The geological time scale at the bottom indicates the major eras, with periods ranging from the Cretaceous to the Quaternary (Q). Plei: Pleistocene Epoch, Plio: Pliocene Epoch. Species distributions: EP (Eastern Pacific), ESP (Eastern South Pacific), ENP (Eastern North Pacific), WA (Western Atlantic), WNA (Western North Atlantic). Photograph credits: image of ESP *H. dipterurus* by F. Zavalaga; image of *H. longus* was taken by L. Bancayán; images of *H. americanus*, *H. sabinus*, and *H. say* were kindly provided by R. Robertson (Robertson and Van Tassell 2023); image of ENP *H. dipterurus* was retrieved from the iNaturalist database (2026). 2009−2024 off−vessel price of *Hypanus brevis*

**Fig. S2.**
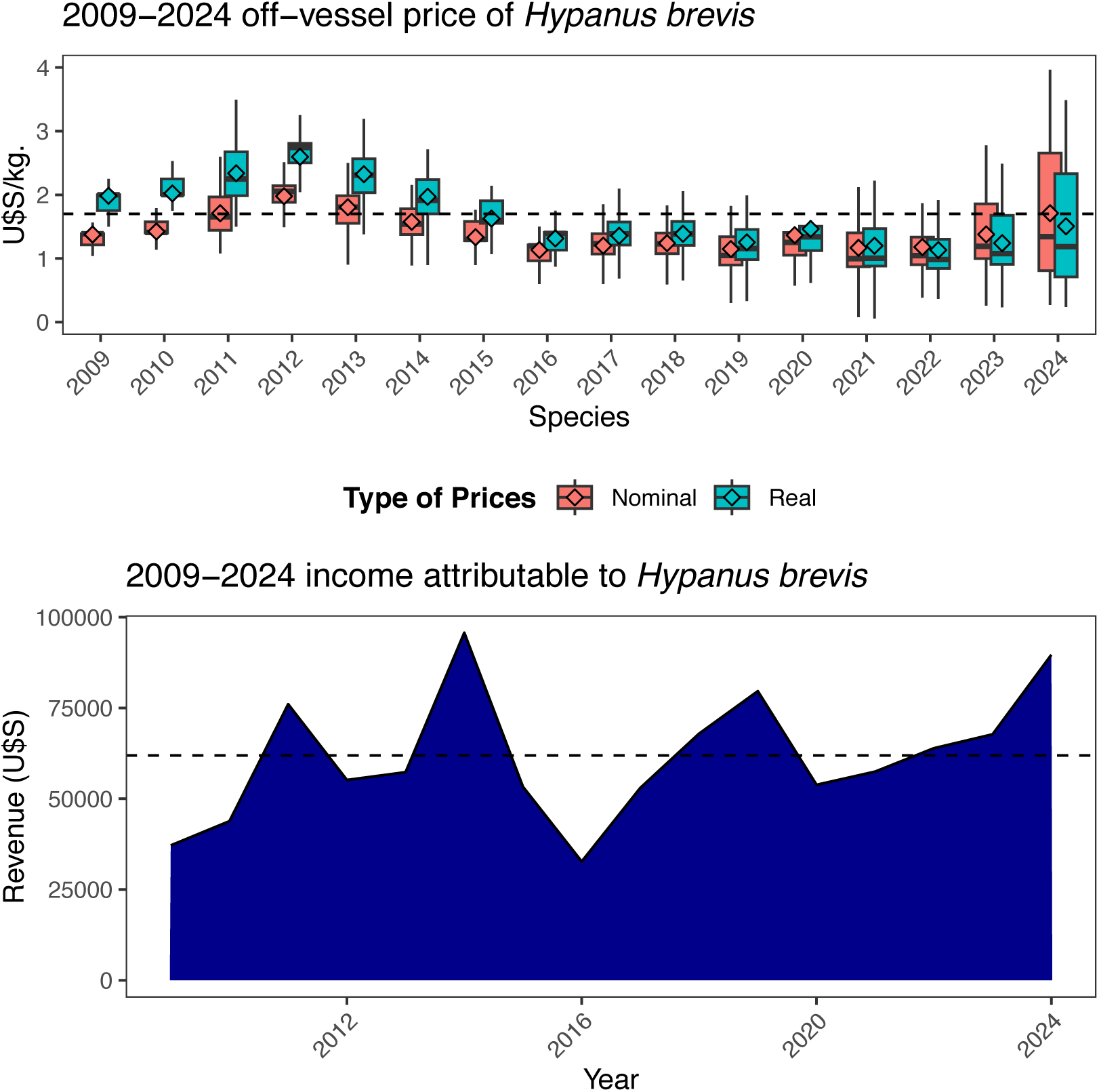
Box-plot with the off-vessel 2009-2024 prices of *Hypanus brevis* (top-panel) and the landing income attributable to this species during the same period (bottom-panel). The horizontal dashed line represents the average 2015-2024 annual income.

**Table S1.**
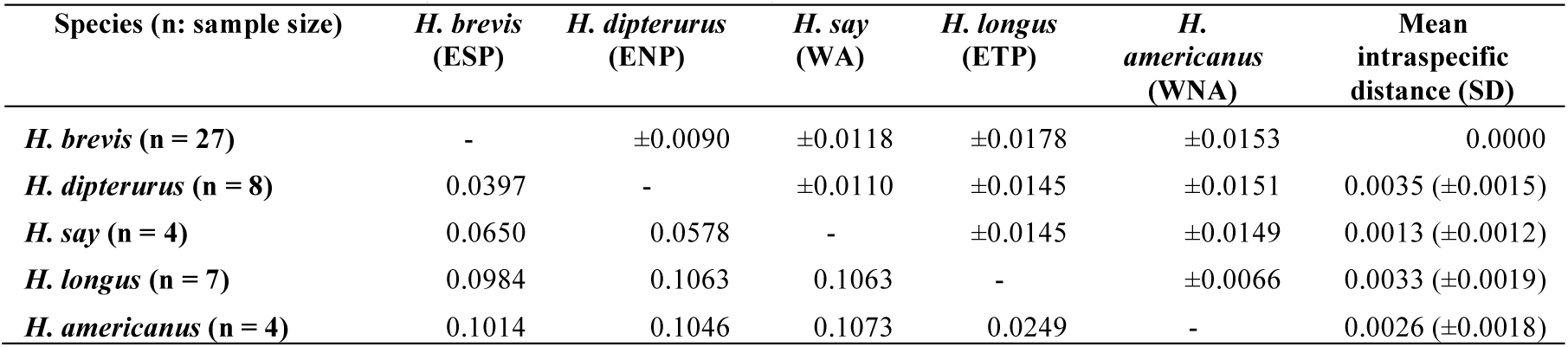
Group interspecific mean distances of a partial COI fragment of five *Hypanus* species from the Pacific (*H. brevis*, *H. dipterurus*, and *H. longus*) and Atlantic Oceans (*H. americanus* and *H. say*). Analyses were conducted using the Kimura 2-parameter model. Interspecific mean distances are shown below the diagonal, while standard error estimates are shown above the diagonal. The last column shows the mean intraspecific distances and their standard error estimates between parenthesis. ESP: Eastern South Pacific; ENP: Eastern North Pacific; WA: Western Atlantic; WNA: Western North Atlantic.

